# Hepatic cytochrome P450 endoplasmic reticulum-associated degradation (ERAD): Topological determinants and cellular partnerships that dictate the preferential P450 proteolytic sorting into macroautophagy rather than UPS

**DOI:** 10.1101/2025.09.25.678692

**Authors:** Xiao Hong, Liang He, Yi Liu, Jinyu Luo, Xiaokun Shu, Maria Almira Correia

**Affiliations:** Departments of Cellular & Molecular Pharmacology, University of California San Francisco, San Francisco CA 94158-2517; Pharmaceutical Chemistry, University of California San Francisco, San Francisco CA 94158-2517; Bioengineering and Therapeutic Sciences, University of California San Francisco, San Francisco CA 94158-2517; The Liver Center, University of California San Francisco, San Francisco CA 94158-2517

**Author notes:** Address correspondence to: M. A. Correia, Mission Bay Campus, Genentech Hall, 600 16th Street, Box 2280, University of California San Francisco, San Francisco, CA 94158-2517. Changping Laboratory, Beijing 102206, China (Xiao Hong).

## Abstract

Many N-terminally endoplasmic reticulum (ER)-anchored cytochrome P450 proteins (P450s) turn over proteolytically via ER-associated degradation (ERAD), others via ER-to-lysosomal-associated degradation (ERLAD), and yet others via both pathways. What precisely dictates their differential proteolytic turnover is unknown. Herein, we employed rabbit liver CYPs 1A1 and 1A2, which reportedly reside in liquid-disorded (l_d_)- and detergent-resistant, liquid-ordered (l_o_)-ER-microdomains, respectively, governed by their specific N-terminal (NT) signal-anchor (SA) subdomains. We now report that this precise SA-dependent ER-topology not only determines the proclivity of CYP1A1 towards ERAD and CYP1A2 towards ERLAD, but also their differential lifespans. We further document that the detergent-resistant l_o_-ER-membranes (DRMs) are morphologically quite similar to mitochondria-associated ER-membranes (MAMs), documented cellular platforms for autophagic-initiation complexes. DRMs and MAMs, composed of saturated fatty-acids, glycosphingolipids and cholesterol, harbor many common morphological markers including the ER-specific prohibitin, erlin-1. Herein employing SURF, a split fluorogenic bifunctional complementation assay, we show that intracellularly, erlin-1 and CYP1A2 interact closely via their ER NT-SAs. siRNA-knockdown (KD) of erlin-1 in HepG2-cells, not only relocated CYP1A2 from DRMs to non-DRMs, but also impaired its ERLAD, resulting in insoluble cellular CYP1A2 aggregates. Upon erlin-1 KD, CYP1A2 ERLAD could be rescued by co-expression of a siRNA-resistant intact erlin-1 or just its NT-1-30 residue SA-domain. Our findings are the first to reveal that the CYP1A2 lifespan and preferential proclivity towards ERLAD is determined by its close association with erlin-1 within DRMs/MAMs. As proof of concept, we document that the ERLAD-proclivity of CYP2B1 is also similarly dependent upon erlin-1-DRM-association.

**SIGNIFICANCE STATEMENT:** The endoplasmic reticulum (ER)-anchored cytochromes P450 (P450s) incur ER-associated degradation (ERAD) and/or ER-to-lysosomal-associated degradation (ERLAD). What determines their preferential proteolytic turnover is unknown. Here, employing P450s CYP1A1 and CYP1A2 that reside in their N-terminal signal anchor-determined liquid-disordered (l_d_)- and detergent-resistant, liquid-ordered (l_o_)-ER-microdomains, respectively, we documented that in HepG2-cells these ER-microdomains determine the CYP1A proteolytic preferences for ERAD vs ERLAD as well as their lifespans. More importantly, we discovered that its intimate interaction with the ER-specific prohibitin erlin-1 colocalized in these l_o_-ER-microdomains, specifically dictates CYP1A2’s ERLAD-preference. Accordingly, siRNA-elicited erlin-1-knockdown disrupted CYP1A2-ERLAD, which was rescued upon coexpression of either a siRNA-resistant erlin-1 or just its N-terminal 1-30 residues. As proof of concept, we document similar characteristics for CYP2B1, another ERLAD-targeted P450.

## INTRODUCTION

The mammalian liver hemoproteins, cytochromes P450 (P450s; MW ≈ 50 kDa) function in the oxidative/reductive metabolism and elimination of numerous endobiotics as well as xenobiotics (pharmacological and recreational drugs, carcinogens, toxins and other foreign substances of dietary or environmental origin) (1, 2). Because of their essential role in the metabolism and elimination of therapeutically relevant drugs, P450s influence not only the efficacy and duration of the pharmacological drug-response, but also the severity and the time course of many pharmacokinetic/pharmacodynamic drug-drug interactions (DDIs) (3–8). Physiological/pharmacological regulation of hepatic P450 content through modulation of its protein synthesis and/or proteolytic turnover as well as its functional alteration through pharmacological inhibition and/or inactivation are thus all therapeutically very relevant issues.

Hepatic P450s are integral endoplasmic reticulum (ER) proteins, exhibiting differential cellular lifespans as well as preferential proteolytic targeting to ER-associated degradation (ERAD) via either ubiquitin (Ub)-mediated 26S proteasomal degradation (UPD) or autophagic-lysosomal degradation (ALD/macroautophagy) (9–12), a process within the ER-to-lysosome-associated degradation (ERLAD) pathway (13–17). However, some P450s may be proteolytically disposed to both cellular degradation pathways, which are complementary in character and exhibit extensive crosstalk with one pathway serving as a failsafe backup for the other under some circumstances (13–17). Despite their common ER-topology, the basis for such preferential P450 proteolytic targeting, is intriguing and unclear.

ER-integral P450s typically share a monotopic, Type 1 topology with the P450 N-terminal (NT) signal anchor (SA) consisting of a ≈ 20-30 residue long amphipatic helix, connected through a linker and a proline-rich segment to the globular P450 catalytic domain, which although embedded in the ER-membrane is largely accessible to the cytoplasm (18–23). A minimal SA length of ≈ 20 residues is considered necessary to span the 30-40 Å-thickness of the ER-bilayer (20). P450 SAs, even among closely related isoforms, are unique in amino acid sequence and serve as the individual P450 signatures (24). Besides serving as P450 ER-anchors, these relatively hydrophobic sequences are also known to oligomerize within the ER-membrane into higher order homomeric/heteromeric complexes, as well as to direct the P450 proteins to relatively dynamic liquid-ordered (l_o_) or liquid-disordered (l_d_) ER-microdomains of defined lipid composition, rather than to a random ER-membrane distribution (25-31; **Fig. 1**). Accordingly, rabbit liver CYPs 1A1 and 1A2 exhibiting ≈ 80% overall sequence identity and 89% sequence similarity, but dissimilar SA sequences, are normally relegated to predominantly l_d_ and l_o_ ER-microdomains, respectively (28–31). CYP2B4, on the other hand, is found in both l_o_ and l_d_-domains, whereas CYP2E1 remains monomeric and is predominantly found in l_d_-domains (29). Furthermore, CYPs 1A1 and 2E1 fully residing in l_d_ ER-microdomains enriched in *unsaturated* fatty acids (FAs), can be readily solubilized by low-strength detergents (i.e.1% Triton X100 or 0.5% Brij 98), whereas the l_o_-residing CYP1A2 is largely resistant to such solubilization and remains firmly lodged in “*detergent-resistant membranes*” or DRMs, enriched in “lipid-rafts” constituted of *saturated* FAs, sphingomyelin (SM), ganglioside GD3 (a glycosphingolipid containing sialic acid and three glycosyl groups) and cholesterol (Chol) (**Fig. 1**; 32-37). Such DRMs can be solubilized by boiling with 8M urea/2M thiourea/4% CHAPS/Tris (TISO) buffer and are distinguishable from non-DRMs by their lower density buoyancy on 5-40% discontinuous sucrose gradients (38–40). Despite the notably low ER-membrane Chol composition (41), methyl-²-cyclodextrin (M²CD)-mediated Chol-depletion of ER-membranes (microsomes) led to the relocation of CYPs 1A2 and 2B4 from l_o_-to l_d_-microdomains, revealing the critical importance of Chol, despite its low ER-content, in the organization of ER l_o_-microdomains or “lipid rafts” (29).

**Fig. 1.**
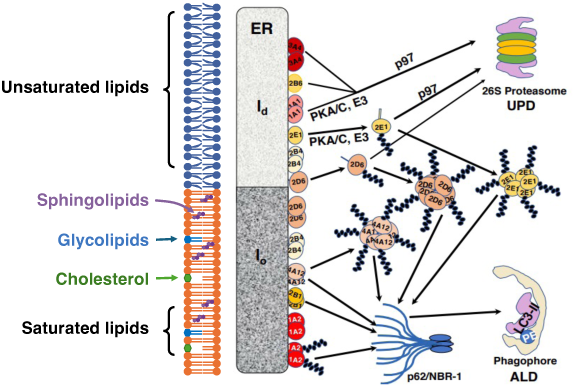
Relative P450 sorting into UPD versus ALD from their localization in l_d_-vs l_o_-ER-microdomains: A schematic representation. While the l_d_-ER-microdomain is enriched in unsaturated fatty acids, the l_o_-ER-microdomain not only is composed of saturated fatty acids but also enriched in cholesterol, sphingolipids and glycolipids. P450s localized to the l_d_-ER-microdomain are extracted from the ER and ferried over by the heterotrimeric p97/Ufd1/Npl4 complex to the 26S proteasome for proteolytic disposal via ERAD/UPD, whereas those in the l_o_-ER-microdomain such as CYP4A11 involve the autophagic receptor p62/SQSTM-1 to chaperone them to the autophagosome for lysosomal disposal via ERLAD/ALD. Note: Organelles/receptors are not drawn to their actual size.

Our preliminary proteomic analyses of relative P450 content in TISO-solubilized DRMs and 1% Triton-soluble fractions derived from hepatocytes of wild-type (WT) mice or ATG5^-/-^-mice with disrupted ALD upon knockout of the essential autophagic initiation gene ATG5 (42), intriguingly revealed that some P450s such as Cyp1a2 were highly enriched in DRMs from ATG5^-/-^-mouse hepatocytes relative to those in WT-mouse hepatocytes. By contrast, P450s in corresponding 1% Triton-solubilized hepatic “non-DRM” fractions such as the constitutive Cyp2e1 or Cyp2b10 showed no relative enrichment upon ATG5-knockdown (43). This hepatic Cyp1a2-enrichment almost exclusively in DRMs (but not in non-DRMs) upon ALD-disruption provided the first clue that the P450s residing in l_o_-but not in l_d_-ER-microdomains may be predisposed to ALD-disposal. To examine this hypothesis, we employed UPD and ALD inhibitors as diagnostic probes along with cycloheximide (CHX)-chase analyses, to compare the relative proteolytic targeting and degradation time-courses of the highly sequence similar rabbit liver CYP1A1 and CYP1A2 proteins. We found that indeed in HepG2-cells, whereas CYP1A1 was preferentially an ERAD/UPD substrate, CYP1A2 was primarily an ERALD/ALD substrate. Examination of the relative structural relevance of their specific ER-SA-anchoring within DRM- and non-DRM-ER-microdomains verified their unique relevance to their individual lifespans and their differential proteolytic targeting. Employing organellar and autophagic markers, we also examined the topological similarity of l_o_-ER-microdomains (DRMs) to mitochondrial-associated ER-membranes (MAMs), dynamic mitochondria-ER apposition sites (44). Among their manifold documented and/or putative physiological and pathological roles (45–52), MAMs have been proposed as contact sites of autophagic biogenesis (37, 44, 53). Our findings identified the ER-specific prohibitin (PHB) family member erlin-1, an <u>ER</u> <u>li</u>pid raft prote<u>in</u> (38–40), as an important determinant of CYP1A2 DRM-localization and ALD. Erlin-1, first documented to reside in ER-derived DRM/lipid rafts (38), upon induction of cell starvation is also reportedly found within MAMs, wherein it intimately associates with the autophagic initiation AMBRA1/BECN1 complex (53). We found that specific siRNA-elicited erlin-1 knockdown (KD), not only switched CYP1A2-localization from DRMs to non-DRMs, but also disrupted its ALD, leading to its accumulation as relatively insoluble cytoplasmic aggregates that required TISO buffer for their solubilization. More importantly, our findings revealed that upon such siRNA-elicited erlin-1-KD, just its NT (1-30 residues; N1-30) ER-anchor was sufficient to rescue not only the normal CYP1A2 ER DRM-localization, but also its disrupted ALD. Furthermore, co-expression of NT CYP1A2 1-33 residues (N1-33) and N1-30 erlin-1-subdomains tagged with a split fluorogenic protein-protein interaction SURF reporter in U2OS and/or HEK293T cells, verified their direct interaction within the ER. Our collective findings are detailed herein.

## RESULTS

### Differential proteolytic targeting of CYPs 1A in HepG2 cells is determined by their individual SA-anchor structure

Given our preliminary proteomic findings of relative Cyp1a2-stabilization in mouse ATG5^-/-^-hepatocytes (43), and that reportedly, rabbit liver CYP1A1 and CYP1A2 reside in different ER-microdomains (28–30), we probed whether this differential ER-topology had any influence on their proteolytic targeting. We therefore examined their individual lifespans and their degradation pathway preferences in HepG2 cells using CHX-chase analyses coupled with diagnostic inhibitors of UPD [Bortezomib, BTZ (54)] and ALD [3-Methyladenine, 3-MA (55) coupled with NH_4_Cl, a lysosomal acidifier) as probes. To readily monitor the proteolytic course of these proteins we also appended a C-terminal (CT) m-Cherry tag, which is photostable upon lysosomal acidification (**Fig. 2 A-F**). We found that indeed while CYP1A1-mCherry was targeted to UPD, as documented by its stabilization by concomitant BTZ-treatment, CYP1A2-mCherry was stabilized by ALD-inhibitors 3-MA/NH_4_Cl and targeted to ALD. Moreover, consistent with its UPD-targeting, CYP1A1 exhibited a shorter half-life (t_1/2_ = 5.93 ± 0.25 h), whereas CYP1A2 also consistent with its ALD-targeting had a relatively longer half-life (t_1/2_ = 12.25 ± 2.87h) in HepG2-cells (**Fig. 2A-F**). This differential ER-microdomain CYP1A localization was reportedly governed by their specific N-terminal (NT)-subdomain and could be switched upon exchange of one CYP1A NT-subdomain with the other (30). To determine whether a similar CYP1A NT-subdomain swap would also influence their relative proteolytic preference, we generated the following NT-CYP1A chimeras: (i) The CYP1A1 NT-1-109 residues were swapped with the corresponding CYP1A2-NT-residues to yield the 1A1(1–109)/CYP1A2 chimeric protein; (ii) the NT-1-107 or 1-205 residues of CYP1A2 were swapped with the corresponding CYP1A1-NT-residues to yield the 1A2(1–107)/CYP1A1 and 1A2(1–205)/CYP1A1 chimeric proteins. Such chimeras have been documented to retain their catalytic function upon co-expression of the electron donor P450-reductase in HEK293T cells, and thus to preserve their structural conformation and functional interactions with their redox partner cytochrome P450 oxidoreductase (CPR) (30). As with their parent, WT CYPs 1A, these proteins were also tagged with a CT-m-Cherry tag. To verify that the m-Cherry tag did not alter the ER-microdomain localization and behaved equivalently to the untagged versions, we similarly employed a simple 1%Triton X-100 solubility assay (30). In this assay, 48 h upon plasmid transfection of m-Cherry-tagged CYP1A proteins or their corresponding NT-chimeras, HepG2 cells were collected and treated with 1%-Triton. The Triton-treated cells were then ultracentrifuged at 100,000 x*g* for 1 h at 4°C with the supernatant (S) fractions containing the Triton-solubilized CYP1A proteins, and the pellets (P) containing the detergent-resistant CYP1A proteins (**Fig. S1A**). Upon pellet-resuspension, equivalent protein amounts of S- and P-subfractions were immunoblotted with an anti-mCherry antibody to determine their relative localization. Consistent with the previous reports of GFP-tagged CYPs 1A (30), CYP1A1-mCherry was largely found in the S-fraction, whereas CYP1A2-mCherry was found in both S- and P-fractions, with a slight enrichment in the latter. Appendage of CYP1A1 NT(1–109) residues onto CYP1A2 changed its distribution in favor of the S-subfraction. Swapping of just CYP1A2 NT-1-107 residues with those of CYP1A1-mCherry led to an equivalent distribution of the 1A2(1–107)/CYP1A1-mCherry chimera in both the S- and P- subfractions, whereas swapping of CYP1A2 NT-1-205 residues with those of CYP1A1-mCherry substantially increased its distribution in the P-subfraction over that of the S-subfraction (**Fig. S1A)**. These findings not only confirmed previous findings (30) that the CYP1A NTs determine their relative ER-microdomain localization, but also verified that this CYP1A behavior was not altered upon incorporation of the mCherry tag.

**Fig. 2.**
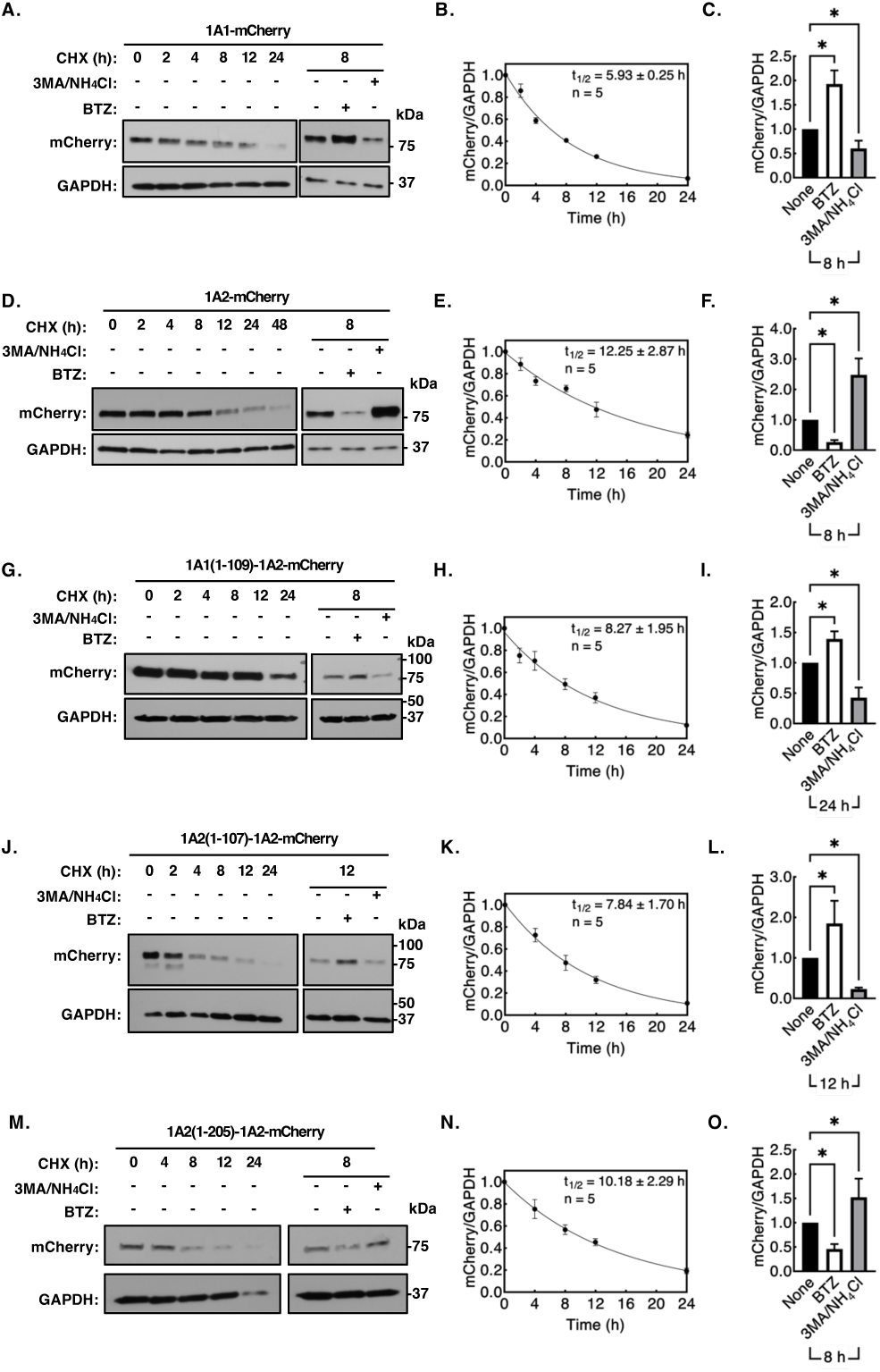
Relative UPD vs ALD proteolytic preferences of the mCherry-tagged parent rabbit liver CYPs 1A and their N-terminal structural chimeras and relative lifespans. A-F. Relative UPD vs ALD proteolytic preferences of the mCherry-tagged parent CYPs 1A1 and 1A2 as probed with CHX-chase analyses and diagnostic UPD (BTZ) and ALD (3MA/NH4Cl) inhibitors. HepG2 cells were seeded in two 6-wells culture plates, and then each well was transfected with 2 μg CYP1A1-mCherry plasmid (**A**) or CYP1A2-mCherry plasmid (**D**). After a 48 h-transfection, HepG2 cells were treated with CHX (50 µg/ml) for indicated times, ± BTZ (10 μM) or ± 3-MA (5 mM)/NH_4_Cl (30 mM) for 8 h. Cells were harvested at indicated times after CHX-treatment and lysates (10 µg) were subjected to IB analyses with GAPDH as the loading control. Densitometrically quantified CYP1A1-mCherry (**B**) or CYP1A2-mCherry (**E**) levels normalized to their corresponding individual GAPDH levels at each harvest time point were quantified relative to their 0 h levels. Values from five experimental replicates were analyzed by Prism Graphpad version 9.5.0 to determine the lifespan [half-life (t_1/2_); mean ± SD] of each mCherry-tagged protein based on a single exponential fit of the data. Upon CHX-chase ± BTZ, or ± 3-MA/NH_4_Cl at 8 h, densitometrically quantified CYP1A1-mCherry (**C**) or CYP1A2-mCherry (**F**) levels normalized to their corresponding individual GAPDH levels and quantified relative to their BTZ or 3-MA/NH_4_Cl levels at 8 h. Values were expressed as Mean ± SD from three experimental replicates relative to their 0 h levels, and the statistical significance was calculated by an ordinary one-way ANOVA analyses. Identical analyses and quantification were conducted upon plasmid transfection and CHX-chase in HepG2 cells of the N-terminal mCherry-tagged chimeras: 1A1(1–109)1A2-mCherry (**G**-**I**), 1A2(1–107)-1A1-mCherry (**J**-**L**) and 1A2(1–205)-1A1-mCherry (**M**-**O**).

To examine the relative ER-l_o_-/l_d_-residency of the mCherry-tagged parent CYPs 1A and their chimeras, we verified their relative fractionation upon a 5-40% discontinuous sucrose gradient ultracentrifugation (**Fig. S1B**). WT CYP1A1 consistent with its (non-DRM) l_d_-residency partitions at the lower higher density levels of the sucrose gradient, whereas WT CYP1A2 is more buoyant and generally partitions at the top, less dense levels of the gradient, consistent with its l_o_/DRM-residency (29, 30). We also found that both mCherry-tagged parent CYP1A proteins partitioned similarly to the corresponding GFP-tagged parent proteins (30), revealing no influence of the C-terminal mCherry tag on their relative partitioning (**Fig. S1B**). On the other hand, the 1A1(1–109)/CYP1A2 chimeric protein exhibited a sucrose-density gradient partitioning similarly to that of the parent CYP1A1 protein, thus revealing that replacing these CYP1A2-NT-residues with the CYP1A1 NT-1-109 residues was sufficient to switch CYP1A2 partitioning, consistent with its findings in **Fig. S1A**. By contrast, whereas the 1A2(1–107)/CYP1A1 chimeric protein exhibited an intermediate distribution between that of its CYP1A1- and CYP1A2-partitioning, the 1A2(1–205)/CYP1A1 chimeric protein was able to acquire more of a CYP1A2-like distribution. This revealed that a much longer CYP1A2 NT-subdomain was necessary to convert the normal CYP1A1 “S” vs “P”-partitioning into that of a CYP1A2-like protein.

Furthermore, CHX-chase analyses coupled with UPD- and ALD-inhibitor probes (**Fig. 2A-O**), also revealed that along with its largely non-DRM/ER l_d_-residency, the 1A2(1–107)/CYP1A1 chimeric protein exhibited only a slightly prolonged lifespan (t_1/2_ ≈ 7.84 ± 1.70 h) relative to that of its parent CYP1A1 protein (t_1/2_ ≈ 5.93 ± 0.25 h) (**Fig. 2J-L**). And consistent with its unchanged proclivity towards UPD, it was also stabilized by the UPD-inhibitor BTZ. By contrast, the 1A2(1–205)/CYP1A1 chimeric protein had a much longer lifespan (t_1/2_ ≈ 10.18 ± 2.29 h) relative to that of its parent CYP1A1 protein (t_1/2_ ≈ 5.93 ±0.25 h) and thus, closer to the lifespan of full-length (FL) CYP1A2 (t_1/2_ ≈ 12.25 ± 2.87h h) (**Fig. 2M-O**). Moreover, in common with CYP1A2, it was also stabilized by the dual ALD-inhibitors 3MA/NH_4_Cl (**Fig. 2M-O**). On the other hand, the 1A1(1–109)/CYP1A2 chimeric protein consistent with its shift to non-DRM/ER l_d_-residency, not only exhibited a slightly shorter lifespan (t_1/2_ ≈ 8.27 ± 1.95 h) relative to that of its parent CYP1A2 protein (t_1/2_ ≈ 12.25 ± 2.87 h), but also switched its proteolytic preference from ALD to UPD (**Fig. 2G-I**). Collectively, these findings revealed that the CYP1A1 NT-subdomain grafted onto CYP1A2 is important not only in determining its ER l_d_-vs l_o_-residency, but also its lifespan. It was also responsible for switching the specific inherent proclivity of CYP1A2 from ALD to UPD. By contrast, a bit longer (1-205 residues) CYP1A2 NT-subdomain grafted onto CYP1A1 was required to switch the inherent l_d_-vs l_o_-residency, lifespan and proclivity of this CYP1A1 chimera towards ALD to more closely resemble that of the parent CYP1A2.

Because just the CYP1A1- and CYP1A2-NT-SAs were found sufficient to determine their specific l_d_-versus l_o_/DRM-residency (30), we probed whether these CYP1A SAs were similarly also sufficient to confer their specific UPD-vs ALD-preference. We therefore examined the proteolytic susceptibilities of just the NT-1-33 residues of C-terminally mCherry-tagged CYP1A1 and CYP1A2 relative to that of their corresponding C-terminally mCherry-tagged parent CYPs 1A (**Fig. 3**). Consistent with the previous report (30) that just the CYP1A NT-(1–33) residue subdomains exhibited comparable ER-targeting profiles to those of their parent CYP1A proteins, we found that indeed they were also sufficient to confer their specific UPD-versus ALD-targeting properties. The finding that the C-terminally mCherry-tagged, NT-(1–33)-CYP1A2 domain still retained its ALD-preference, suggests that despite the multiple ubiquitinatable K-residues and phosphorylatable S/T-residues present on the m-Cherry-tag external surface that might have influenced it towards UPD, the degradation of this chimera was still predisposed towards ALD, and thus, unaltered from that of its parent FL CYP1A2 (**Fig. 3**).

**Fig. 3.**
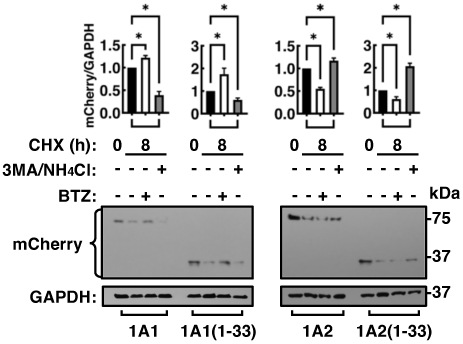
Relative UPD vs ALD proteolytic preferences of the full-length (FL) parent rabbit liver CYPs 1A C-terminally mCherry-tagged and their corresponding similarly mCherry-tagged NT-1-33-residue-structural subdomains. HepG2 cells were seeded into four 6-well culture plates, plasmids (2 μg DNA) for FL 1A1-mCherry, 1A1(1–33)-mCherry, FL 1A2-mCherry, and 1A2(1–33)-mCherry were transfected into each well. After 48 h transfection, HepG2 cells were subjected to CHX-chase ± BTZ, or ± 3-MA/NH_4_Cl for 8 h, and analyzed similarly to CYP1A1-mCherry (**C**) or CYP1A2-mCherry (**F**) (Fig. 2). Values were expressed as Mean ± SD from three experimental replicates relative to their 0 h levels, and the statistical significance was calculated by an ordinary one-way ANOVA analysis. All analyses were performed by Prism Graphpad version 9.5.0. Data indicate as mean ± SD of N = 3. *Note the NT-subdomains retain the proteolytic preferences of their parent CYP1A proteins*.

#### The proteins associated with isolated DRM fractions are identical to those known to associate with MAMs

Consistent with our preliminary proteomic findings of Cyp1a2-enrichment in DRM-fractions of ATG5^-/-^-mouse hepatocytes (43), and its partitioning in the upper less-dense sucrose-gradient fractions (**Fig. S1B**), our preliminary immunoblotting (IB) analyses revealed that the Cyp1A2 in these sucrose fractions was also associated with the ALD-essential protein ATG5 (*not shown*). Intriguingly, ATG5 (as well as ATG14L, and other ALD-associated ATG-proteins) reportedly also associate with mitochondria-associated ER-membranes (MAMs), which are documented to constitute autophagosomal initiation sites (37, 44, 53). Notably, the composition of such mitochondrial and ER-contact sites closely resembles that of DRMs in harboring lipid-raft like microdomains enriched in cholesterol, glycosphingolipids, saturated fatty acids, etc, in addition to certain specialized ER-specific PHB-like erlin-1 and −2 proteins (38, 39, 53). To determine whether the CYP1A2-associated DRMs isolated by our sucrose gradient fractionation resembled MAMs, we isolated DRMs via a 5-40% discontinuous sucrose-density gradient centrifugation (**Fig. 4A)** in parallel with a MAM-isolation from the same CYP1A2-transfected HepG2-cell culture (**Fig. 4B**). We then probed our isolated DRM- and MAM-fractions for various established MAM organellar markers (37–39, 53) such as: Erlin-1 (MAM), mitofusin 2 (MFN2, a mitochondrial protein that bridges MAM mitochondria and ER-contact sites), P450-reductase (ER), Vdac1 (a mitochondrial and MAM protein) and GAPDH (a soluble cytoplasmic marker protein) (**Fig. 4A** & **B**). As in **Fig. S1B**, CYP1A2 distribution, consistent with its lipid raft/DRM association was the highest in sucrose fractions 4-6. These CYP1A2-enriched fractions were pooled together as DRMs, whereas fractions 8-10, wherein CYP1A1 usually partitions (**Fig. S1B**) were considered as non-DRMs. Each pooled fraction was then probed by IB-analyses for ATG5 and the above organellar markers. Consistent with its 1%Triton-soluble non-DRM characteristics, cytoplasmic GAPDH was associated largely with the non-DRMs (**Fig. 4A**). By contrast, the MAM markers erli-1, VDAC1 and MFN2 were largely associated with the DRM-fractions (**Fig. 4A**). The parallel MAM isolation from the same CYP1A2-transfected cells yielded MAMs as well as MAMs tightly associated with mitochondria, along with cytosol and ER (**Fig. 4B**). When probed with corresponding organellar marker antibodies, MAMs and MAM/Mitochondria fractions were shown to be largely associated with MFN2, ATG5, and erlin-1, whereas the ER and cytoplasmic 1% Triton-soluble fractions were associated with P450 reductase (CPR) and GAPDH, respectively. The findings of erlin-1, ATG5, VDAC1 and MFN2 in both CYP1A2-comprised DRMs and MAMs thus establish that ER-DRMs and cellular MAMs indeed share common morphological markers and are quite similar (**Fig. 4**).

**Fig. 4.**
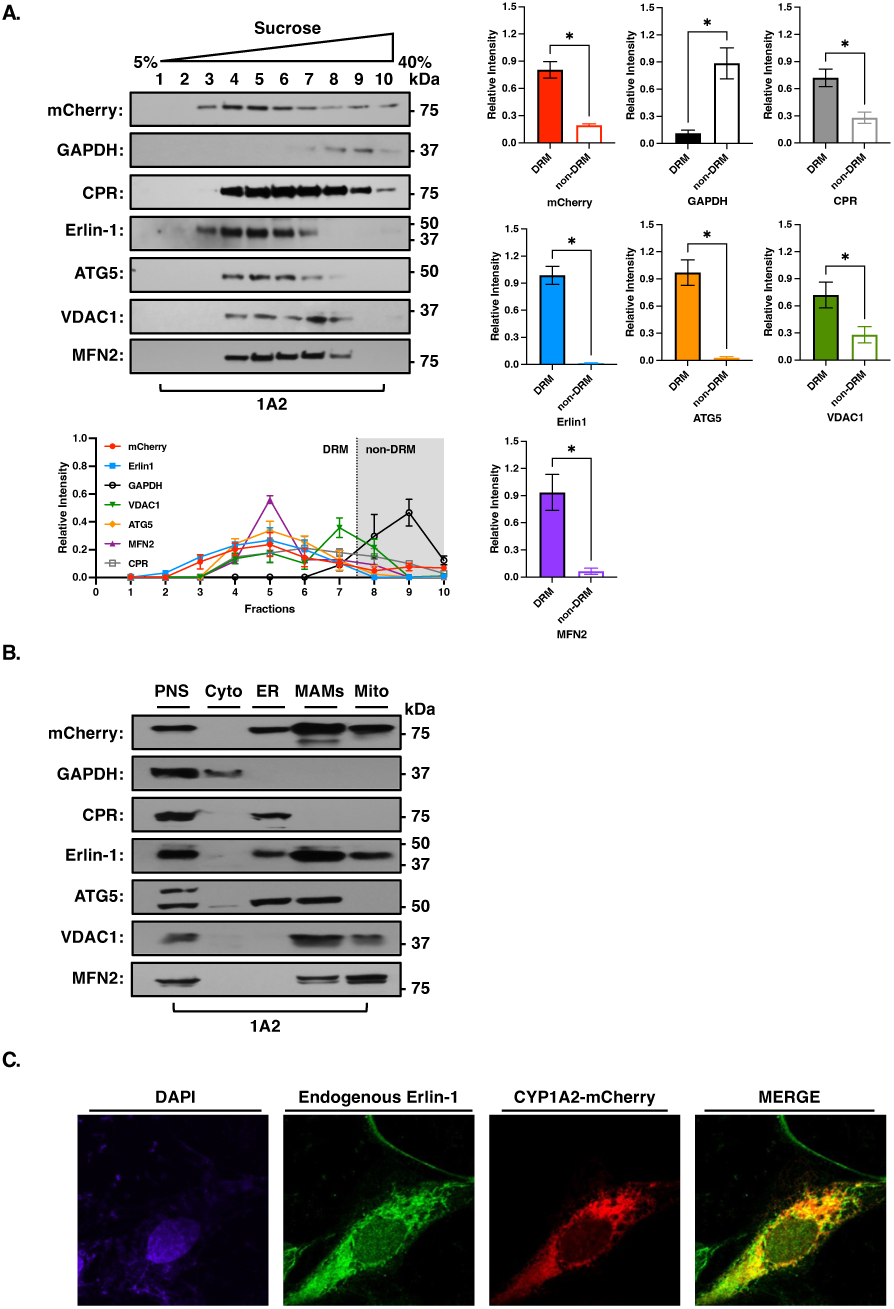
Relative subcellular fractionation of FL CYP1A2-mCherry into ER-DRMs and cellular MAMs and IB analyses of their commonly shared morphological markers. Two subcellular fractionation procedures were carried out in parallel employing cultured HepG2 cells expressing CYP1A2-mCherry and split into two batches. **A**. Cells in one batch were treated with 1% v/v Triton X-100 on ice for 30 min. A linear 5-40% sucrose density gradient method was employed to isolate DRMs. Gradient fractions (1 mL) each were carefully collected starting from the top of the gradient. **B.** Mitochondria-associated ER-membranes (MAMs) were isolated as per the protocol detailed in Materials and Methods. The isolated fractions from each procedure were then subjected to Western IB analyses. The blots were probed with antibodies specific to mCherry, glyceraldehyde-3-phosphate dehydrogenase (GAPDH), autophagy-related protein 5 (ATG5), cytochrome P450 reductase (CPR), ER lipid raft-associated protein 1 (erlin-1), voltage-dependent anion channel 1 (VDAC1), and mitofusin 2 (MFN2). *Note that CYP1A2 in the upper buoyant sucrose (DRM) fractions is associated with the ALD-essential protein ATG5 (A), and ER-DRMs and cellular MAMs share common morphological markers such as erlin-1 and MFN2 (B)*. **C**. ER-colocalization of both erlin-1 and CYP1A2 in U2OS cells by confocal microscopic analyses. For experimental details see Materials & Methods. Endogenous erlin-1 within the cells was detected immunochemically with a rabbit erlin-1 primary polyclonal antibody followed by a secondary antibody consisting of goat anti-rabbit IgG Alexa Fluor 488, which rendered it visible as green fluorescence. The overexpressed CYP1A2-mCherry was visualized as red fluorescence and the colocalization of erlin-1 and mCherry was documented upon merging of their respective fluorescence channels.

Confocal microscopic analyses of U2OS cells transfected with a CYP1A2-mCherry plasmid, fixed, permeabilized and after blocking with normal goat serum, stained overnight first with erlin-1 polyclonal antibody, and then by goat anti-rabbit IgG Alexa Fluor 488 (**Fig. 4C**), verified the intimate ER-colocalization of both erlin-1 and CYP1A2, plausibly in common ER-lipid-raft domains. CYP1A2 was also found to exhibit a patchy, grainy appearance, resembling autophagic puncta/aggregates that did not quite colocalize with erlin-1 (**Fig. 4C**). Because CYP1A2-mCherry was overexpressed, this CYP1A2-aggregate accumulation may reflect the relative excess of CYP1A2-mCherry over the endogenous erlin-1 cellular content.

#### CYP1A2 and erlin-1 cellular interactions: A role for erlin-1 in CYP1A2-ALD?

The NT-regions (NTs; residues N1-30) of erlins-1 and −2 are responsible for their specific targeting to ER-lipid-raft microdomains (38–40). Given the involvement of MAMs and erlin-1 in autophagic initiation (37, 44, 53), and the immunochemically detectable co-localization of CYP1A2 and erlin-1 in ER DRM/MAM lipid-rafts (**Fig. 4C**), we next considered whether any direct interactions of CYP1A2 NT-sequences (NT1-33 residues) with those of erlins-1 and −2 would be plausible. Clustal Omega multiple sequence analyses revealed that CYP1A2 had a higher number of identical/similar residues to erlin-1 NT and erlin-2 NT than CYP1A1 (**Fig. S2**). Given the NT-sequence identity/similarity between the two erlins and CYP1A2, and the fact that the NTs of many P450s including CYP1A2, are known to homo- and heterodimerize with each other or other co-resident proteins such as their electron-donor P450-reductase (CPR) within the ER-membrane bilayer (25–28), we considered the possibility that erlin-1 and CYP1A2 NTs could also similarly interact and heterodimerize within the DRM/MAM ER-microdomains in the course of CYP1A2-ALD. To verify this possibility, we employed the split fluorogenic SURF-bimolecular functional complementation assay, wherein we engineered vectors for each protein tagged at its CT-domain with a split fluorescent SURF-reporter (n- and c-UnaG-tags) (55). Should the two proteins indeed intimately interact intracellularly, then they would fluoresce upon reconstitution of the intact SURF-reporter. Our confocal microscopic analyses of U2OS cells co-expressing plasmids with split n-SURF and c-SURF tags revealed that upon co-expression, full-length (FL) erlin-1-nSURF indeed interacted with the FL parent CYP1A2-c-SURF (**Fig. 5A**), as did erlin-1-NT (N1-30)-nSURF with CYP1A2-NT (N1-33)-cSURF (**Fig. 5C**). Similar interactions of full-length erlin-1-nSURF with the parent CYP1A2-c-SURF were also observed upon co-expression in HEK293T cells (**Fig. S3**). As expected, CYP1A2-NT (N1-33)-nSURF and CYP1A2-NT (N1-33)-cSURF also cross interacted (i.e. homodimerized) (**Fig. 5B**), as did the parental CYP1A2-nSURF and CYP1A2-cSURF upon co-expression in HEK293T cells (**Fig. S3**). Furthermore, that this interaction was specific and not due to any spurious interactions, was verified by the lack of any fluorescent interactions upon confocal analyses of either U2OS (**Fig. 5D**) or HEK293T cells (**Fig. S3)** co-transfected with ER-anchored erlin-1-nSURF and cytoplasmic c-SURF-tagged ornithine decarboxylase 1 (ODC), findings that confirmed their different cellular locations and served as a negative control. By contrast, although FL erlin-1 and FL erlin-2 interacted as expected upon co-transfection of HEK293T cells (**Fig. S3**), no comparable interactions were detected upon cotransfection of erlin-1-NT (1–30)-nSURF and FL erlin-2-cSURF (**Fig. S3**). This is consistent with the fact that the erlin1/2-dimerization domain lies beyond their NT-1-30 residues (38–40). It is also notable that in U2OS cells while both FL erlin-1 and CYP1A2 proteins strictly localized in the ER and perinuclearly, their NT-subdomains upon expression were found to be distributed throughout the entire cells, including the nuclei (**Fig. S7**), plausibly reflecting the absence of any intrinsic nuclear export sequences in their structure.

**Fig. 5.**
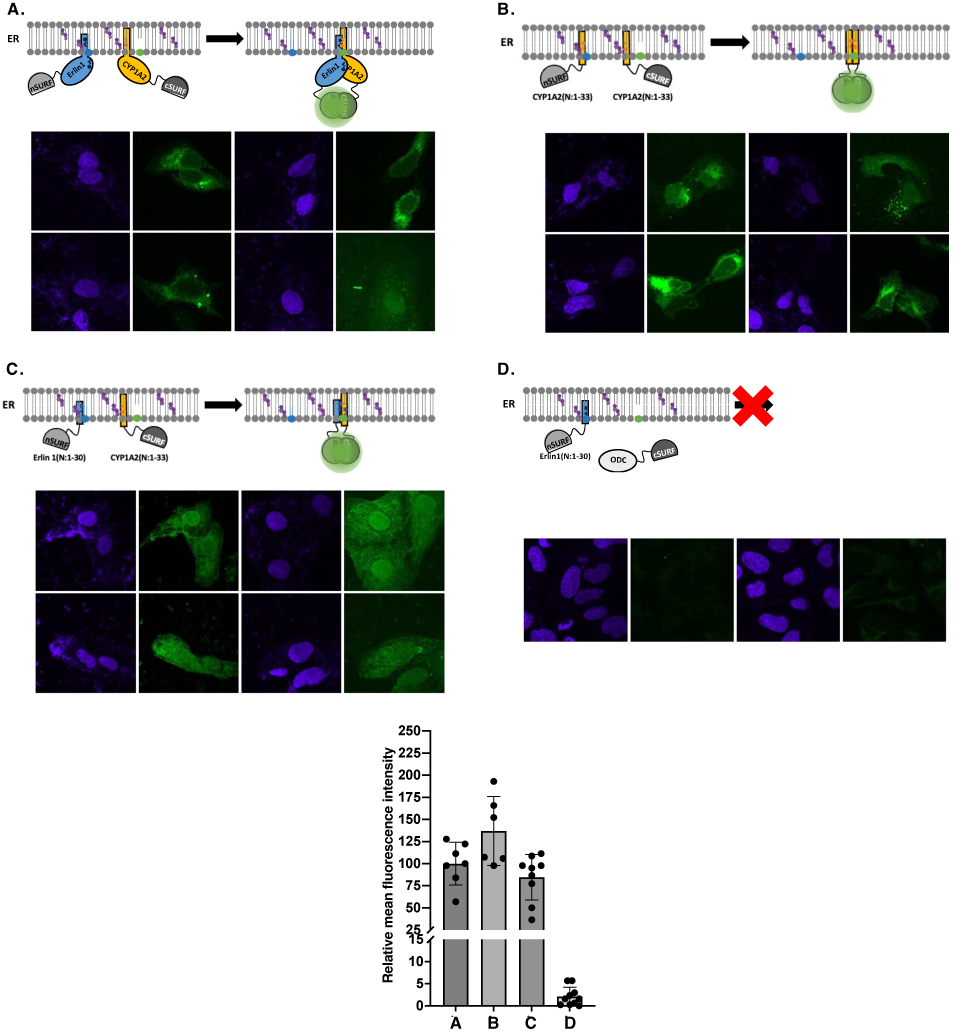
Split fluorogenic SURF-bimolecular functional complementation assays document that CYP1A2- and erlin-1-NT-SA subdomains are sufficient for their fluorescent interactions within the ER-microdomains. Confocal microscopic analyses of U2OS cells co-transfected with: FL erlin-1-nSURF and FL CYP1A2-cSURF revealing fluorescent interactions of the two proteins (**A**). N(1–33)-CYP1A2-nSURF and N(1–33)-CYP1A2-cSURF subdomain interactions document the homodimerization of the CYP1A2 SAs within the ER (**B**). N(1–30)-erlin-1-nSURF and N(1–33)-CYP1A2-cSURF subdomain fluorescent interactions establish that these NT-subdomains are sufficient for their interactions within the ER (**C**). A negative assay control documenting that no fluorescent interactions are detected between FL ER-anchored erlin-1-nSURF and cytoplasmic ODC-cSURF (**D**), thereby excluding any spurious interactions in **A**-**C**. *Note that unlike the FL-proteins which are largely retained within the ER (A), the NT-subdomains exhibit intranuclear import (B, C)*.

To probe whether erlin-1 plays any role in CYP1A2 colocalization in ER lipid-rafts/DRMs, we examined the effect of erlin-1 siRNA-knockdown (KD) on CYP1A2-DRM fractionation upon 5-40% discontinuous sucrose gradient sedimentation, after preliminary optimization of erlin-1 KD temporal course (**Fig. S4**). Erlin-1-KD directed CYP1A2 from the more buoyant DRM-fractions to the lower density non-DRM sucrose gradient-fractions (**Fig. 6A**). To further examine whether in fact erlin-1 is involved in CYP1A2-ALD, we monitored the effects of its siRNA-KD on CYP1A1 and CYP1A2 degradation through CHX-chase analyses at the 8 h-time point. The scrambled siRNA as the corresponding parallel control failed to alter CYP1A1 or CYP1A2 degradation or their individual preference for UPD or ALD, respectively, as documented with the specific diagnostic probes, BTZ or 3-MA/NH_4_Cl of each degradation process (**Fig. 6B**). By contrast, erlin-1 siRNA-KD, considerably mitigated the RIPA-buffer solubilizable CYP1A2-fraction that is normally stabilized by 3-MA/NH_4_Cl (Soluble; *compare lanes 12 vs 16*), and to a much lesser extent also some of that of CYP1A1-fraction that is committed to ALD (Soluble; *compare lanes 4 vs 8*) (**Fig. 6B**). To determine whether this CYP1A-reduction reflected enhanced loss either due to irreversible proteolysis or RIPA-insoluble protein aggregation, we solubilized the RIPA-insoluble lysate fractions (pellets) of CYP1A1 and CYP1A2 from both siRNA-KD and control siRNA-treated HepG2 cells with TISO-buffer (**Fig. 6B**). Notably, very little CYP1A1 was found in TISO-solubilized CYP1A1 fractions from control siRNA-treated cells at 0 h, and this further decreased at 8 h of CHX-chase, irrespective of UPD- or ALD-inhibition (*Pellet; lanes 1*-*4*). Upon erlin-1 siRNA-KD, the levels of TISO-solubilized CYP1A1 noticeably increased, and were appreciable following ALD inhibition at 8 h of CHX-chase (**Fig. 6B**; *Pellet; compare lanes 5 and 8*), suggesting that the degradation of the minor CYP1A1 fraction normally committed to ALD was impaired upon erlin-1 KD. By contrast, the TISO-solubilized CYP1A2 fraction was appreciable at 0 h, and greatly increased upon ALD-inhibition at 8 h of CHX-chase in control siRNA-treated cells (**Fig. 6B**; *Pellet; compare lanes 9 and 12*). Even more remarkably, upon erlin-1 siRNA-KD, all TISO-solubilized CYP1A2 fractions substantially increased in size, both at 0 h and 8 h of CHX-chase. Thus, the TISO-solubilized CYP1A2 fraction not only was sizable at 0 h, but also further increased upon ALD-inhibition at 8 h of CHX-chase (**Fig. 6B**; *Pellet*; c*ompare lanes 9 vs 12 and 13 vs 16, and particularly lanes 12 vs 16*). These findings indicated that upon erlin-1 siRNA-KD, the apparent loss of RIPA-solubilized CYP1A2 fraction observed upon ALD-inhibition was not due to proteolytic loss as it could be recovered as TISO-solubilizable aggregates (**Fig. 6B**; *Compare lanes 12 in RIPA-soluble versus TISO-soluble pellet fractions*). Together, these findings revealed that erlin-1-KD impaired CYP1A2 ALD, thus indicating a plausible role for erlin-1 in CYP1A2-ALD. This plausible role of erlin-1 in CYP1A2-ALD could be due to its known involvement in recruitment of the autophagosome initiation complex AMBRA1/BECN1 (37, 53) to MAMs, and/or simply due to its buttressing heterodimeric interactions with its ER-co-resident CYP1A2 within DRM/lipid raft/MAM during ALD.

**Fig. 6.**
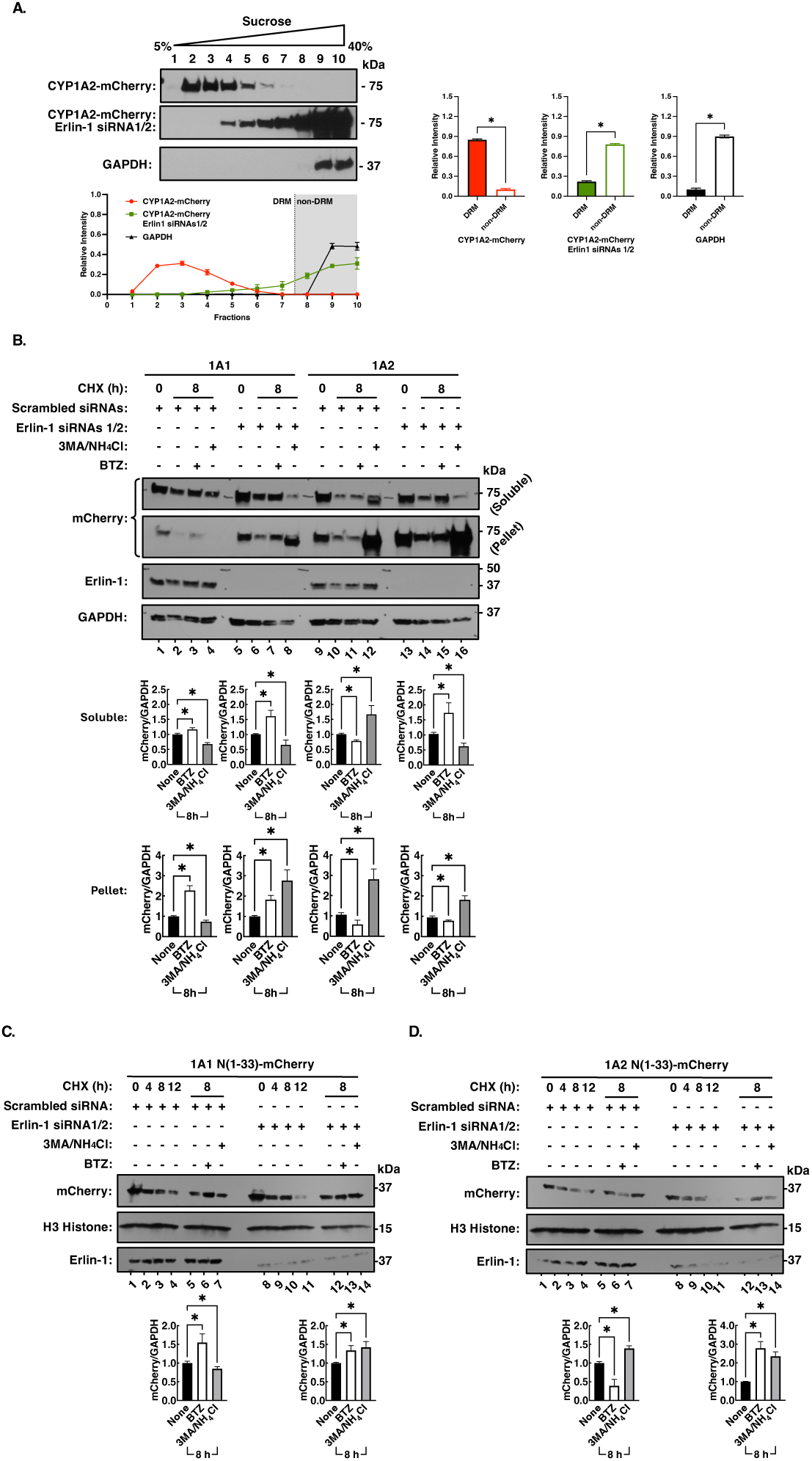
**Erlin-1 siRNA-KD reverses CYP1A2 ER-topology from DRMs to non-DRMs and impairs CYP1A2-ALD, resulting in CYP1A2-aggregate build-up. Similar impairment of CYP1A2**(**1–33**)**-ALD upon erlin-1 siRNA-KD.** HepG2 cells were transfected with the CYP1A2-mCherry plasmid. After 12 h, the culture medium was replaced with fresh MEM containing either scrambled siRNAs (control) or erlin-1 siRNA1 plus siRNA2 (erlin siRNA1/2) for erlin-1 KD. After an additional 24 h, the medium was replaced with fresh MEM containing either the scrambled or erlin-1 1/2 siRNAs, and the cells were incubated for another 24 h before being harvested. The samples were lysed in hypotonic buffer and sonicated at low speed (600 g) to remove nuclei and cell debris. **A**. The lysates were then incubated in a membrane solubilization buffer containing 1% Triton X-100, cleared and subjected to a 5-40% sucrose gradient subfractionation by ultracentrifugation (210,000 g/19h) for DRM isolation. **B**. HepG2 cells were transfected with the CYP1A2-mCherry plasmid and treated with the scrambled or erlin-1 1/2 siRNAs exactly as described above. Cells were processed, and the 1% Triton-treated cell lysates were subjected to a 5-40% sucrose gradient subfractionation as described above. Subfractions 7-11 were collected, diluted with 1.15 M KCl and resedimented by ultracentrifugation at 105,000*g*/1 h. The pellet was collected, resuspended and designated as the “soluble” fraction. Subfractions 4-6 were also collected, diluted with 1.15 M KCl and resedimented by ultracentrifugation at 105,000*g*/1 h, The pellet was resuspended in TISO buffer and designated as the “pellet” fraction (see Materials & Methods for further experimental details). HepG2 cells were similarly transfected with plasmids for CYP1A1(1–33)-mCherry (**C**) or CYP1A2(1–33)-mCherry (**D**). Transfected cells were then treated with scrambled siRNAs or erlin-1 1/2 siRNAs for erlin-1 KD and processed exactly as described above. After 48 h of siRNA-treatment, CHX was added at 0 h and the degradation of CYP1A1 NT(1 33)-mCherry and CYP1A2 NT(1–33)-mCherry subdomains was monitored through CHX-chase analyses from 0-12 h and the effect of diagnostic UPD (BTZ, 10 μM) and ALD [3-MA (5 mM)/ NH_4_Cl (30 mM)] inhibitors monitored at 8 h. Cell lysates (10 μg) were subjected to IB analyses with H3-Histone as the loading control.

We therefore further explored the effects of erlin-1 siRNA-KD or control siRNA on the relative degradation of CYP1A1 NT (1–33)-mCherry and CYP1A2 NT (1–33)-mCherry subdomains through CHX-chase analyses (**Fig. 6C**). As with their corresponding FL parent mCherry-tagged CYPs 1A (**Fig. 6**), control siRNA failed to alter CYP1A1 NT (1–33)-mCherry and CYP1A2 NT (1–33)-mCherry-degradation and/or their individual preference for UPD or ALD, respectively (**Figs. 3** and **6C**). However, by contrast and similarly to that of its parent CYP1A2-protein, erlin-1 siRNA-KD not only considerably reduced the CYP1A2 NT (1–33)-mCherry fraction that is normally destined to ALD and/or is stabilized by 3-MA/NH_4_Cl, but also to a lesser extent increased the relative CYP1A1 NT(1–33)-mCherry-fraction that is normally committed to ALD (**Fig. 6C**; *CYP1A2*(*1–33*)*-mCherry compare lanes 7 &14; CYP1A1*(*1–33*)*-mCherry compare lanes 7 &14)*.

To further define the role of erlin-1 NT in CYP1A2-ALD, we co-transfected FL CYP1A2-mCherry-Myc-His expression plasmid with the following plasmids: A control (scrambled) erlin-1 siRNA plasmid, an erlin-1 siRNA-KD plasmid, and/or either an erlin-1 site-directed mutant plasmid or a non-mutated erlin-1 NT (1–30)-mCherry plasmid (**Fig. 7**). The erlin-1 mutant plasmid while retaining the native erlin-1 amino acid sequence is resistant to the erlin-1 siRNA-KD documented above due to its mutated DNA nucleotide composition (**Fig. 7**). Consistent with the above observations (**Fig. 6A**), erlin-1 siRNA-KD changed the DRM-localization of CYP1A2 to that of non-DRM, whereas control scrambled siRNAs had no such effect (**Fig. 7A**). Co-transfection of erlin-1 siRNA-KD along with the siRNA-resistant mutant erlin-1 plasmid to a large extent preserved the DRM-status of CYP1A2, and more surprisingly, so did the co-transfection of just the erlin-1 NT (1–30)-mCherry expression plasmid (**Fig. 7A**). In parallel, when the effect of these erlin-1 siRNAs was examined on CYP1A2 degradation, control siRNAs were unable to affect the profiles of CYP1A2-ALD or its 3MA/NH_4_Cl-elicited inhibition at 8 h of CHX-chase (*lanes 1-4*), whereas (as in **Fig**. **6**), erlin-1 siRNA-elicited KD reduced the 3MA/NH_4_Cl-inhibitable CYP1A2-fraction normally committed to ALD (**Fig. 7B**; *lanes 6-8*). Co-transfection of erlin-1 siRNA-KD along with the siRNA-resistant erlin-1 mutant expression plasmid yielded a very similar profile of CYP1A2-degradation as that with the control siRNAs, thereby validating that the mutant erlin-1 was indeed resistant to siRNA-KD and functioned equivalently to the WT cellular erlin-1 in restoring the CYP1A2-ALD profile (**Fig. 7B).** When just the mCherry-tagged erlin-1 NT (1–30) plasmid which is inherently siRNA-resistant due to its intrinsic nucleotide sequence was co-expressed, it could by itself similarly rescue CYP1A2-ALD characteristics. These findings thus documented that the erlin-1 NT (1–30) is not only as efficient as the WT erlin-1, but also is sufficient to restore CYP1A2-ALD upon siRNA-KD of the native cellular erlin-1 (**Fig. 7**).

**Fig. 7.**
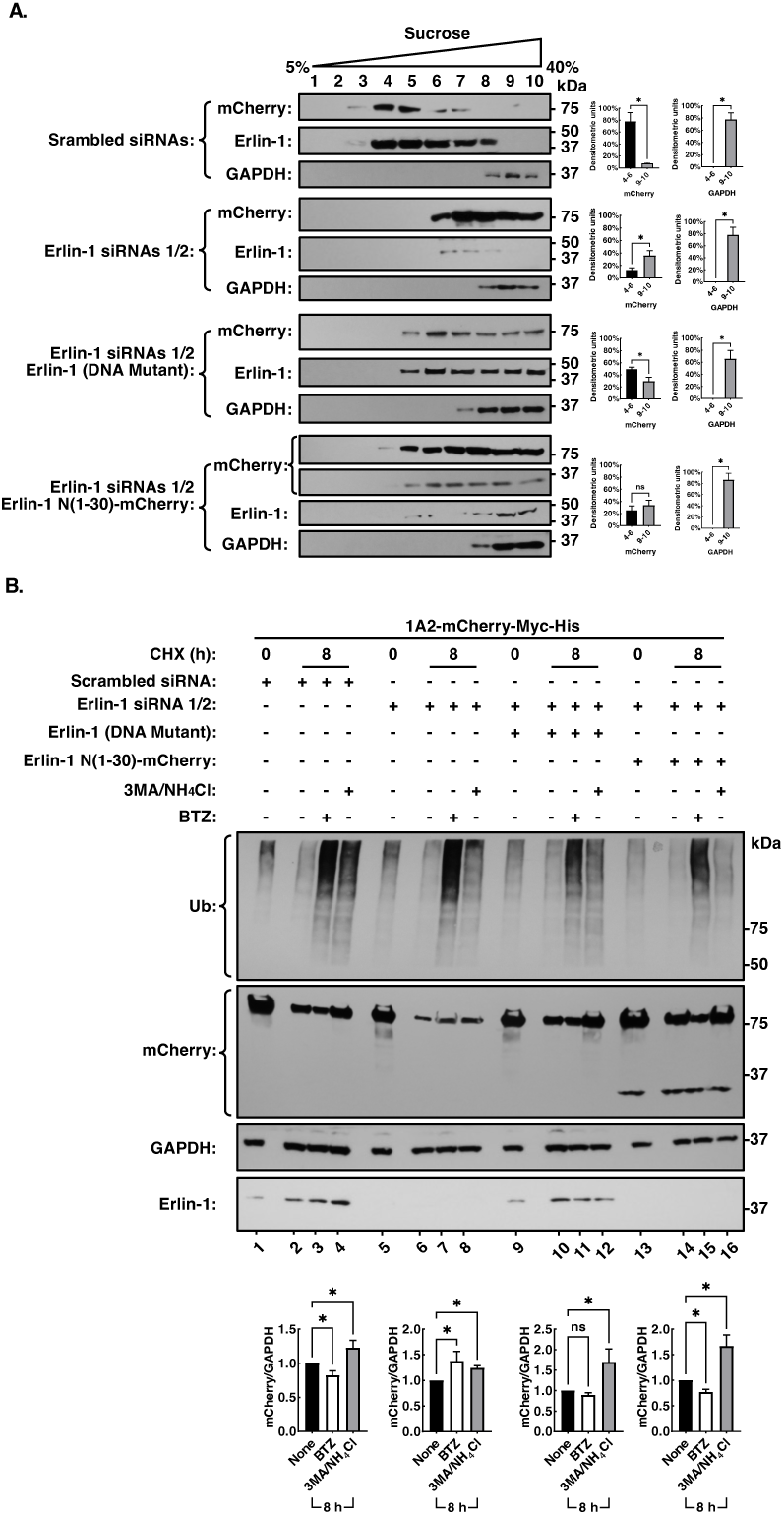
Co-expression of siRNA-resistant erlin-1 or even just its NT. (**1–30**) **subdomain is sufficient to reverse the loss of CYP1A2-mCherry DRM-buoyancy and CYP1A2-mCherry ALD upon siRNA-elicited erlin-1-KD.** HepG2 cells were transfected with the CYP1A2-mCherry plasmid exactly as described in Fig. 6. After 12 h, the culture medium was replaced with fresh MEM containing the scrambled siRNA (control) or siRNAs 1/2. Twelve h after siRNA treatment, the medium was replaced again, and fresh scrambled siRNA (control) or siRNAs 1/2 plasmids along with either erlin-1 (DNA mutant) or erlin-1 (1–30)-mCherry plasmid was added and the cells incubated for another 24 h. The medium was again replaced with fresh MEM containing the same siRNAs, as well as erlin-1 (DNA mutant) or erlin-1 (1–30)-mCherry plasmids and the cells were cultured for an additional 24 h. **A**. To determine the relative subfractionation of CYP1A2 in DRM-vs non-DRM-subfractions, cleared cell lysates were subjected to a discontinuous 5-40% sucrose-gradient ultracentrifugation exactly as described (Fig. 4). Gradient fractions (1 mL) each were carefully collected starting from the top of the gradient and aliquots of these gradient subfractions subjected to IB analyses with mCherry-IgGs (CYP1A2) and GAPDH-IgG (controls). Note that the loss of CYP1A2-DRM localization upon erlin-1 siRNA, is reversed upon co-expression of the siRNA resistant erlin-1 DNA mutant plasmid or even just the erlin-1 N(1–30) subdomain. B. Cells were treated with scrambled siRNA (control) or erlin-1 1/2 siRNAs, and subsequently with either erlin-1 (DNA mutant) or erlin-1 (1–30)-mCherry plasmid, exactly as described above. After the second 24 h-incubation period, cells were treated with CHX (50 μg/mL) at 0 h, with or without BTZ (10 µM) or 3-MA (5 mM)/NH_4_Cl (30 mM) for 8 h. Cell lysates (10 µg) were subjected to mCherry or Ub-IB analyses, with GAPDH as a control. *Note that the loss of CYP1A2-ALD upon erlin-1 siRNA 1/2-KD (Compare lanes 8 vs 4), is reversed upon coexpression of either erlin-1 DNA-mutant (Compare lanes 12 vs 8) or just erlin-1 N*(*1–30*) *subdomain (Compare lanes 16 vs 8)*.

#### Proof of concept: Differential proteolytic degradation of P450s of the CYP2B and CYP4A subfamilies

As proof of concept of our hypothesis based on the above findings that the differential CYP1A ER-microdomain topology is a determinant of their corresponding proteolytic pathway preferences, we examined two different sets of P450 proteins pertaining to subfamilies 2B and 4A. Despite their at the least 75% amino acid sequence identity, and >85% sequence similarity as well as the highly similar tertiary fold of the crystallized mammalian hepatic CYPs 2B (57–59), we and others have previously documented that the proteolytic degradation of rat liver CYP2B1 and human liver CYP2B6 preferentially occurs via ALD and UPD, respectively (60–62). Thus, we selected this P450 pair to test our hypothesis that the differential proteolytic targeting of CYPs 2B1 and 2B6 proteins would similarly also be conferred by differences in their ER-microdomain topology. Consistent with our existing literature evidence (12, 60–62), CHX-chase analyses confirmed that in HepG2 cells, CYP2B1 also exhibited not only its proteolytic preference for 3MA/NH_4_Cl-inhibitable ALD but also consistently had a longer lifespan (t_1/2_ ≈ 10.35 ± 2.81 h) than CYP2B6 (t_1/2_ ≈ 6.87 ± 1.79 h), a confirmed BTZ-inhibitable UPD substrate (**Fig. 8A, B**). We also verified that as in the case of CYP1A NTs, this preference was similarly conferred by their NT-(1-33 residues) SAs (**Fig. S5**). A simple 1% Triton X100-solubility test upon centrifugal fractionation also verified that although both proteins were found in the 1% Triton X100-soluble “S” and 1% Triton X100-insoluble “P” subfractions, CYP2B1 partitioned to a greater extent in the P vs S-subfraction, whereas quite the reverse was true of CYP2B6 (**Fig. S5**). Nonlinear sucrose (5-40%) gradient fractionation revealed that similarly to CYP1A2, CYP2B1 partitioned in the lower density, higher buoyancy “DRM”-subfractions 3-6, whereas in common with CYP1A1, CYP2B6 was found in the higher density “non-DRM”-subfractions 7-9 (**Fig. 8C, D**). The CYP2B1-enriched DRM-subfractions were also associated with the ALD-initiating ATG5 enzyme and the MAM-markers erlin-1 and VDAC1, as well as its electron donor CPR (**Fig8C, D**). A direct comparison of CYP1A2 and CYP2B1 MAMs from HepG2-cells subjected to erlin-1 siRNA-interference, revealed that similarly to CYP1A2-siRNA-KD findings, along with their erlin-1 content, CYP2B1 content of MAMs was greatly reduced upon erlin-1 siRNA-KD relative to that in corresponding cells treated with the scrambled siRNAs (**Fig. 8E**). These findings thus confirmed the very similar DRM-lipid raft ER-microdomain topology and erlin-1-dependence of CYP2B1 to that of CYP1A2, along with their common proteolytic ALD preference.

**Fig. 8.**
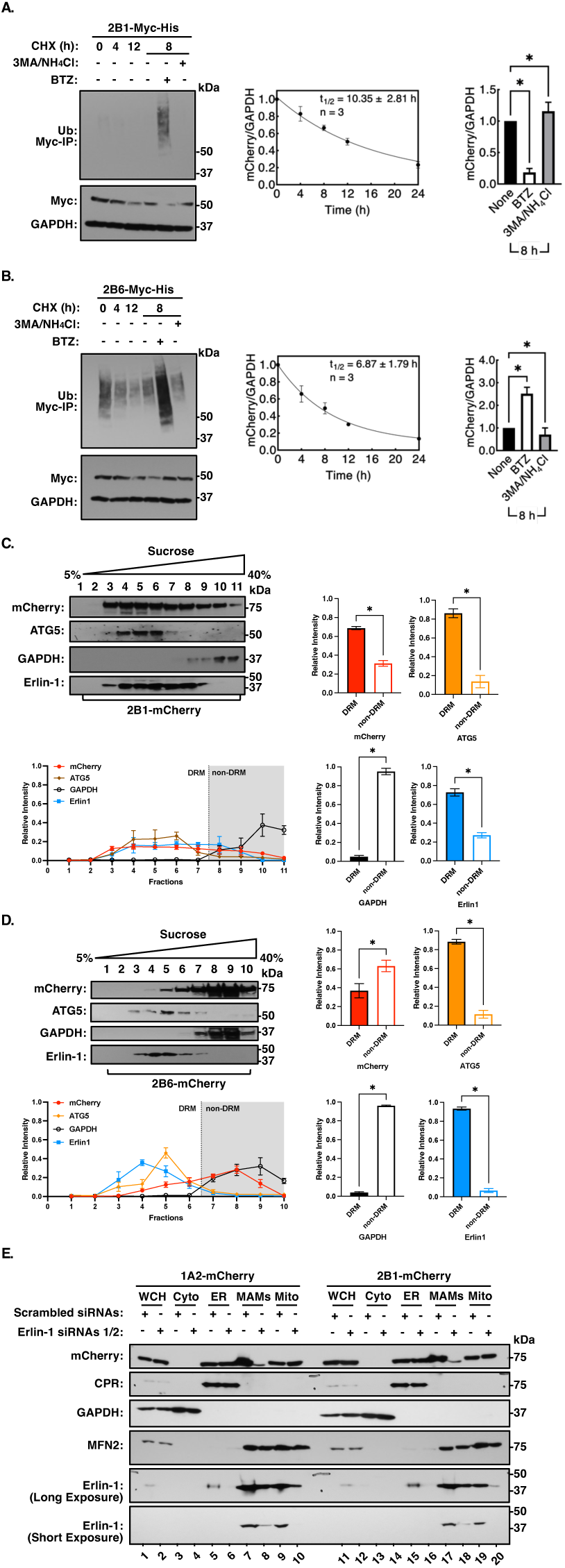
Proteolytic turnover of rat hepatic CYP2B1 and human hepatic CYP2B6 as proof of concept: Similarities between CYP1A2 and CYP2B1. A. The two highly related P450 subfamily 2B proteins are known to exhibit differential proteolytic preferences for UPD and ALD as verified with the diagnostic UPD (BTZ) and ALD (3-MA/NH_4_Cl) inhibitor probes as well as life-spans consistent with these proteolytic preferences. **B**. Nonlinear 5-40% sucrose density gradient ultracentrifugal subfractionation reveals ALD-prone CYP2B1 to reside in the buoyant DRM-fractions, whereas UPD-prone CYP2B6 is found in the more sucrose dense non-DRM fractions. As expected, CYP2B1 in DRM-subfractions is also associated with ATG5, erlin-1 and VDAC1. CYP2B6 in non-DRM-subfractions is not. **C**. Erlin-1 siRNA-KD reveals that CYP2B1 consistent with the findings of CYP1A2, also relies on erlin-1 for its localization in the DRM/MAM-fractions. Except for the use of CYP2B1 and CYP2B6 plasmids, other experimental details are exactly as provided in Figs. 2 and 6 as well as Materials & Methods.

The second P450 protein set we tested was based on our very recent report (63, 64) that P450s of the CYP4A subfamily, despite their very high primary sequence relatedness (77% sequence identity and 93% similarity), also exhibited differential proteolytic preferences. Thus, human liver CYP4A11 and mouse liver Cyp4a12a are targeted preferentially to ALD, in contrast to mouse liver Cyp4a10 that is targeted largely to UPD. We thus subjected CYP4A11 and Cyp4a10 to the simple 1%-Triton X100-solubility test by centrifugal fractionation as well as DRM-isolation by the nonlinear 5-40% sucrose density fractionation (**Fig. S6**). In common with CYPs 1A and CYPs 2B (**Fig. S6**), we found that although both CYP4A11 and Cyp4a10 proteins were found both in the 1% Triton X100-soluble “S”- and 1% Triton X100-insoluble “P”-subfractions, that CYP4A11 partitioned to a relatively greater extent in the “P”-vs “S”-subfraction, whereas Cyp4a10 partitioned to a greater extent in the “S”-rather than “P”-subfraction (**Fig. S6B**), consistent with their specific proteolytic preferences. Moreover, the nonlinear sucrose density fractionation, also relegated CYP4A11 to the lower density, more buoyant DRM-subfractions 4-6, along with ATG5, the MAM markers erlin-1, MFN2 and VDAC1 (**Fig. S6C**). Collectively, these findings confirm our hypothesis that the DRM/lipid raft/MAM ER-microdomain topology of CYPs 1A2, 2B1 and plausibly 4A11, not only is erlin-1-dependent, and conferred by their corresponding NT-SAs, but also determines their proteolytic preferences for specific targeting to the ALD versus UPD pathway.

## DISCUSSION

Our findings described above confirmed previous reports that CYP1A1 and CYP1A2 exhibit differential ER-topology, residing in l_d_- and l_o_-ER-microdomains (the latter a.k.a ER-lipid-rafts/DRMs), respectively, which is to a large extent dictated by their individual NT-SAs. We now document that this CYP1A NT-SA-dependent ER-microdomain topology also dictates the preferential proteolytic turnover of CYP1A1 via ERAD/UPD and CYP1A2 via ERLAD/ALD, and consequently determines their relative lifespans. Replacing one CYP1A NT-subdomain containing its SA for the other, not only switched their corresponding l_d_- and l_o_-ER-microdomain topologies, but also altered their individual normal lifespans and preferential proteolytic targeting. Furthermore, a C-terminally mCherry-tagged CYP1A1 or CYP1A2 SA-subdomain was sufficient to confer this proteolytic preference. More importantly, we also documented that such preferential proteolytic targeting of CYP1A2 into ERLAD/ALD is also dependent on its colocalization with erlin-1 in these l_o_-ER-lipid-raft-microdomains.

Erlins-1 and −2 are ER-specific prohibitins (PHBs), N-terminally integrated into the ER-lipid-raft microdomains. They are known to homo- and hetero-oligomerize as well as to form large multimeric complexes (38–40), although their interacting/dimerizing subdomains are thought to reside outside their ER-anchoring NTs. Intriguingly, the vast majority (>70%; 28, 29) of hepatic CYP1A2 is also found to be clustered within the l_o_ ER-lipid-rafts/DRMs, wherein their selective clustering may contribute not only to their reported catalytic/functional interactions (28, 65), but also to their segregation as autophagic cargo. Herein, we documented that indeed siRNA-elicited erlin-1-KD not only led to the redistribution of CYP1A2 from buoyant DRMs to 100,000*g*-sedimentable non-DRM-fractions (**Figs. 6A** & **7A**), but also to a dramatic reduction in the CYP1A2-fraction normally committed to ALD that is fully assessable upon 3MA/NH_4_Cl-ALD inhibition, along with a relative increase in its normally small, UPD-committed BTZ-inhibitable fraction (**Fig. 6B**). Remarkably, along with CYP1A2 ALD-impairment, such erlin-1-KD also led to a marked accumulation of insoluble cytoplasmic CYP1A2 aggregates that required high urea (8 M)/thiourea (2M)/CHAPS-containing TISO buffers for their solubilization (**Fig. 6B**). Intriguingly, upon siRNA-KD of native erlin-1, co-expression of just its NT (1-30 residues) was sufficient to restore CYP1A2 to its normal DRM-localization (**Fig. 7**), thus implicating a critical role for the erlin-1 ER-anchor in CYP1A2-ALD. Our SURF-bimolecular functional complementation assay coupled with confocal microscopic analyses indeed revealed that CYP1A2-NT(1–33) ER-anchor does heterodimerize with the erlin-1 NT(1–30) ER-anchor (**Fig. 5**), and thus may well be responsible for retaining CYP1A2 in the ER-lipid-rafts/DRMs/MAMs. Optimal cellular CYP1A2 ALD apparently critically relies on a well-balanced, cellular erlin-1 level proportionate to that of CYP1A2. When upon erlin-1-KD, cellular CYP1A2 levels exceed those of erlin-1, CYP1A2 ALD is impaired, resulting in correspondingly increased insoluble CYP1A2-aggregates (**Fig. 6B**). Similarly, when upon exogenous CYP1A2-overexpression, the CYP1A2 level exceeds that of the endogenous erlin-1 basal content, patchy granular CYP1A2-clumps were detected upon confocal microscopic analyses (**Fig. 4C**), even in the absence of any obvious autophagic disruption. Although erlin-1 and erlin-2 are known to heterodimerize and function jointly (38–40), and our confocal studies revealed that CYP1A2 also interacts with erlin-2 (**Fig. S3**), the precise extent of erlin-2’s plausible contribution to CYP1A2 ALD remains to be determined. Collectively, these findings argue that erlin-1 may play a critical dual complementary role in CYP1A2 ALD: (i) by fostering the proper normal incorporation and/or retention of soluble CYP1A2 into ER-lipid rafts, thereby enabling CYP1A2 clustering as autophagic cargo. And (ii) given its crucial role as an integral component of ER-lipid rafts, by ensuring the structural integrity of these functional platforms for the recruitment of the autophagic initiation AMBRA1/BECN1 and Atg5-ATG12-ATG16L-complexes required for autophagosomal biogenesis.

Intriguingly, the true organellar source of the isolation membrane or phagophore that gives rise to the nascent autophagosome upon autophagic induction has been controversial. In recent years, cumulative literature evidence from several different research teams converges on the mitochondria-associated ER membranes or MAMs as a, if not, the site of autophagosomal biogenesis (37, 44, 53). Such ER-mitochondria contact sites are thought to be tethered together by the mitochondrial protein mitofusin 2 (MFN2), and to contain dynamic ER-lipid rafts composed not only of ER-specific PHB-like proteins erlins-1 and −2, but also sphingoglycolipids such as the ganglioside GD3, cholesterol and calnexin (CANX) as well as the mitochondrial voltage-dependent anion channel 1 (VDAC1). MAMs have been shown to serve a functional platform for the recruitment of autophagic initiation complexes such as ATG14L, ATG5, AMBRA/BECN1, etc (44, 53). Upon amino acid starvation-elicited autophagic induction, the earliest key events in autophagosomal biogenesis reportedly entail the molecular association of CANX with the ganglioside GD3, AMBRA 1 and WIPI1 (37, 53). Accordingly, siRNA-mediated KD of MFN2, of the GD3-synthetase ST8SIA1, or of erlin-1, each impaired not only the enhanced molecular association of CANX-AMBRA1 normally observed upon autophagic induction, but also the detection of autophagic LC3 puncta (37), revealing the critical importance of undisrupted ER-lipid rafts to MAM-elicited recruitment of core molecular autophagic initiator complexes.

Our IB analyses of CYP1A2-containing ER-lipid-rafts/DRMs and MAMs isolated in parallel from the same CYP1A2-plasmid transfected HepG2 cell cultures, revealed in addition to CYP1A2, the presence of many common constituents such as MFN2, VDAC1, erlin-1 and associated autophagic initiation factors (ATG5 and/or its covalent ATG5/ATG12 complex, detectable as a 50 kDa species). These common constituents reinforce the notion that MAMs may indeed represent hubs for autophagic initiation of CYP1A2 ALD. Notably, given that nearly 25% of the ER lipid-bilayer consists of phosphatidylethanolamine (PE) almost exclusively on its outer leaflet (66), the recruitment of the autophagic initiation Atg5-ATG12-ATG16L complex to these ER-lipid raft-microdomains would greatly facilitate its catalytic covalent PE-conjugation to LC3-I to generate the hallmark autophagosomal morphological marker, LC3-II (67). Subsequent elongation and LC3-II-mediated recruitment of autophagic receptors such as SQSTM1/p62 and/or NBR1 and their associated protein cargos would lead to the autophagosomal maturation that progresses to protein ALD.

Although our studies have not specifically examined the role of any of these ER-lipid raft/DRM- and MAM-lipid constituents in CYP1A2-ALD, other reports have amply documented the critical importance of the GD3-ganglioside for starvation-induced autophagic initiation through GD3-synthetase ST8SIA1-KD (37). Similarly, despite the relatively low physiological ER-Chol content, M²CD-mediated Chol-depletion of ER-membranes is shown to lead to the CYP1A2-relocalization from l_o_ ER-lipid rafts/DRMs to l_d_ ER-nonDRMs (29, 30). Our analyses of the CYP1A1(1–109)/CYP1A2 chimera have shown that its relocalization to l_d_ ER-nonDRMs led not only to its proteolytic switch from ERLAD/ALD to ERAD/UPD, but also to the shortening of the lifespan of this chimeric CYP1A1(1–109)/CYP1A2 protein relative to that of its parent CYP1A2. Similar M²CD-mediated Chol-depletion of ER-membranes if not reversing the CYP1A2 proteolytic preference, would be expected at the very least to impair CYP1A2-ALD, simply by disrupting the essential functional lipid-raft platform required for the autophagic initiation complex recruitment.

As proof of concept that their specific P450-ER-lipid raft/DRM/MAM-microdomain topology is a critical determinant of their preferential ALD-targeting, we have examined two other P450 protein-pairs from two different CYP2B- and CYP4A-subfamiles: Specifically, rat liver CYP2B1 versus human liver CYP2B6, as well as human liver CYP4A11 versus mouse liver Cyp4a10. These specific P450 pairs in common with CYPs 1A, also exhibit differential proteolytic turnover preferences for ALD and UPD (12, 60–64). In fundamental support of our basic hypothesis, we found that both CYPs 2B1 and 4A11 which are established ALD-substrates, were found to reside in ER-lipid-raft/DRM/MAM-microdomains, in close association with erlin-1 and other MAM-morphological markers. In the case of CYP2B1, we have shown this ER-microdomain localization to be also dependent on erlin-1 (**Fig. 8C**). By contrast, the relatively shorter lifespan proteins CYP2B6 and Cyp4a10 which were found in ER-non-DRM-microdomains were found to be preferential UPD-targets (12, 63, 64. Notably, while the P450 ER lipid-raft/DRM/MAM-microdomain topology is clearly a determinant of the ALD-preference of some hepatic P450s, other plausible mechanistic scenarios may also exist: For instance, although the N-terminally ER-anchored constitutive CYP2E1 has been shown to be normally monomeric, to reside in l_d_-ER-microdomains (29), and to exhibit a short lifespan (≈ 7 h) and a documented UPD-preference (12, 69), in the presence of its inducers i.e. acetone, ethanol, isoniazid, etc) this lifespan is prolonged to ≈ 37 h, and the CYP2E1 protein acquires a proclivity for ALD (61, 68, 69). Intriguingly, whether this is due to its altered switch from its normal ER-localization to l_o_-ER-microdomains, or its known propensity for cellular trafficking between the ER- and non-ER-organelles including the plasma membrane, during which it risks being captured by cytoplasmic autophagic receptors and thus marked as cellular autophagic cargo, remains to be determined. Nevertheless, given their remarkable capacity to transit between erlin-1-dependent lipid raft/DRM- and non-DRM-microdomains within the ER-bilayer, we posit that the ER-anchored P450s are excellent models to probe and unravel the cellular mysteries and intricacies involved in the physiologically relevant ERAD/ERLAD processes.

**MATERIALS & METHODS:**

### Plasmids, vectors, and viruses

Human CYP1A1 and CYP1A2 were amplified by PCR using CYP1A1-GFP and CYP1A2-GFP constructs (30) as templates. The rat CYP2B1 cDNA amplified by PCR with pSW1 encoding the full length rat CYP2B1 as the template (62), and CYP2B6 fragment amplified by PCR from HepG2 cell cDNA were subsequently cloned into the pcDNA3.1-Myc-His(-)-A_CMV-F_H10 vector, as described (12). Erlin-1-FLAG was purchased from Sino Biological, erlin-2-Myc-FLAG was purchased from OriGene, and pET28a-His6-ODC1-Ctag was a gift from Dr. Mark Howarth (Addgene plasmid #163614). The amplified fragments were cloned into pmCherry-N1 Plasmid (Clontech PT3974-5) or pcDNA3.1 vectors to generate the expression plasmids 1A1-mCherry, 1A2-mCherry, 2B1-mCherry, 2B6-mCherry, erlin-1-mCherry, and erlin-2-mCherry.

To explore the domain functions, chimeric constructs were generated by exchanging homologous regions between CYP1A1 and CYP1A2 as previously reported (30). Specifically, the N-terminal (NT) fragment of CYP1A1 (residues 1-109) was used to replace the corresponding region of CYP1A2, yielding 1A1(1–109)1A2-mCherry. Conversely, CYP1A2 fragments (residues 1-107 or 1-205) were exchanged with the corresponding regions of CYP1A1 to generate 1A2(1–107)1A1-mCherry and 1A2(1–205)1A1-mCherry plasmids, respectively. To assess the contribution of the NT signal-anchor sequences, truncation mutants encoding only the NT-residues were cloned in-frame with either mCherry or SURF (the n-SURF and c-SURF plasmids were provided by the Shu lab (56)], resulting in 1A1(N:1-33)-mCherry, 1A2(N:1-33)-mCherry, 1A1(N:1-33)-nSURF, 1A1(N:1-33)-cSURF, 1A2(N:1-33)-nSURF, 1A2(N:1-33)-cSURF, 2B1(N:1-30)-mCherry, and 2B6(N:1-30)-mCherry. Similarly, the first 30 NT-residues of erlin-1 were fused with nSURF or cSURF to generate erlin-1(N:1-30)-nSURF and erlin-1(N:1-30)-cSURF.

For fusion-tag constructs, a C-terminal (CT) Myc-His or Myc-epitope tag was inserted downstream, resulting in 1A1-mCherry-Myc-His, 1A2-mCherry-Myc-His, 1A1(1–109)1A2-mCherry-Myc-His, 1A2(1–107)1A1-mCherry-Myc-His, 1A2(1–205)1A1-mCherry-Myc-His, 2B1-Myc-His, 2B6-Myc-His, 2B1-Myc, 2B6-Myc, 2B1(N:1-30)-mCherry-Myc-His, and 2B6(N:1-30)-mCherry-Myc-His.

In addition, full-length CYP1A1, CYP1A2, erlin-1, erlin-2, and ornithine decarboxylase 1 (ODC1) were fused with either nSURF or cSURF to generate 1A1-nSURF, 1A1-cSURF, 1A2-nSURF, 1A2-cSURF, erlin-1-nSURF, erlin-1-cSURF, erlin-2-nSURF, erlin-2-cSURF, and ODC1-cSURF.

All truncations, domain swaps, and tagged fusion constructs were generated by PCR-based cloning or homologous recombination using the NEBuilder® HiFi DNA Assembly Kit (New England Biolabs, MA) and their authenticity was verified by DNA sequencing. The primers, templates, and vectors used for plasmid construction are summarized in Supporting Information (**Table 1**).

#### *E. coli* transformation and culture

Bacterial transformations were performed using Takara competent cells (Cat. #636763, Takara Bio) according to the manufacturer’s instructions (69). Briefly, competent cells (100 μL) were thawed on ice for 5 min, gently resuspended, and transferred into a 14-mL round-bottom Falcon tube (Cat. #149591B, Fisher Scientific). Plasmid DNA (10 ng) was added, and the mixture was incubated on ice for 30 min followed by heat shock at 42°C for 45 s. The cells were immediately chilled on ice for 5 min and then resuspended in pre-warmed Super optimal medium with catabolic repressor (SOC; Cat. #C994M61, Thomas Scientific) to a final volume of 1 mL. After incubation at 37°C with shaking at 225 rpm for 1 h, kanamycin (Cat. #BP861, Sigma-Aldrich) containing Lysogeny broth (LB) medium (Cat. #L3522, Sigma-Aldrich) was added to bring the culture volume to 4 mL, and cells were further incubated overnight at 37°C with shaking at 225 rpm.

##### HepG2/U2OS cell cultures

Cells were cultured as recently described (63). Briefly, HepG2 cells were cultured in minimal Eagle’s medium (MEM, #10370021; Gibco) containing 10% v/v fetal bovine serum (FBS, #10099-141C; Thermo-Fisher Scientific) and supplemented with nonessential amino acids (#11140050; Thermo-Fisher Scientific) and 50,000 U penicillin-streptomycin (#15140122; Thermo-Fisher Scientific) and 1 mM sodium pyruvate (#11360070; Thermo-Fisher Scientific). U2OS cells were cultured in McCoy’s 5A Modified Medium (#16600108; Gibco), supplemented with 10% FBS and 50,000 U penicillin-streptomycin. All cells were cultured at 37°C under 5%CO_2_/95%O_2_.

##### HepG2 cell transfections

HepG2 cells were seeded 24 h prior to transfection and cultured until ∼70% confluence. DNA-lipid complexes were prepared by diluting plasmid DNA in Opti-MEM reduced-serum medium (Opti-MEM; Cat. #31985062, Thermo-Fisher Scientific) at a ratio of 1 μg plasmid DNA per 100 μL Opti-MEM. TransIT-X2 transfection reagent (Cat. #MIR 6004, Mirus Bio) was then added at 3 μL per 1 μg DNA, and the mixture was gently pipetted to ensure complete mixing. After incubation at room temperature for 20 min, the complexes were added dropwise to the culture wells, followed by gentle rocking of the plate to evenly distribute the transfection complexes. Cells were maintained for 48 h post-transfection. The culture medium was replaced with fresh MEM medium 12 h after transfection and subsequently changed every 24 h.

##### Cycloheximide (CHX)-chase analyses of CYP1A degradation and treatment with diagnostic UPD or ALD probes

HepG2 cells were seeded in 6-well plates at a density of 2 mL MEM medium per well, 24 h prior to transfection and transfected with the indicated plasmids. Forty-eight h post-transfection, cells were treated with CHX (50 μg/mL; Cat. #C801N89, Thomas Scientific) for the indicated time periods to monitor protein degradation. To determine the degradation pathways, cells were exposed to diagnostic UPD or ALD inhibitor probes under the following conditions: (i) untreated control (CHX only, 50 μg/mL); (ii) proteasome inhibition, with CHX 50 μg/mL + bortezomib (BTZ, 10 μM; Cat. #C835F72, Thomas Scientific); and (iii) autophagic–lysosomal inhibition, with CHX 50 μg/mL + 3-methyladenine (3-MA, 5 mM; Cat. #C910A04, Thomas Scientific), and NH_4_Cl (30 mM; Cat. #C988H67, Thomas Scientific). Cells were harvested at the designated time points for subsequent analyses. Each condition was performed in triplicate. Protein degradation was assessed by mCherry immunoblotting (IB) followed by densitometric quantification. Half-life values (t_1/2_) were calculated using one-phase exponential decay fitting in GraphPad Prism 6.07. Data are presented as mean ± standard deviation (SD) of at the least 3 different experiments.

##### Immunoprecipitation (IP) analyses of ubiquitinated proteins

HepG2 cells were collected and washed three times with phosphate-buffered saline (PBS; Cat. #10010002, Gibco) at 4°C by centrifugation at 2,000 × g for 5 min to remove residual culture medium. The cell pellets were resuspended in 100 μL RIPA lysis buffer (pH 7.4) supplemented with protease inhibitor cocktail (Cat. #5871S, Cell Signaling Technology), 1 mM phenylmethylsulfonyl fluoride (PMSF; Cat. #8553S, Cell Signaling Technology), and 10 mM antipain (Cat. #C802G16, Thomas Scientific). Cell suspensions were homogenized on ice and further disrupted using an Omni Sonic Ruptor 250 sonicator at 20% amplitude (50 W) for three cycles of five pulses each, with at least 1 min intervals between cycles. Lysates were clarified by centrifugation at 600 × g for 5 min at 4°C in an Eppendorf Centrifuge 5425 (Cat. #EP5405000441, Sigma-Aldrich) to remove unbroken cells and nuclei, and the supernatants were collected. Protein concentrations were determined using the bicinchoninic acid (BCA) assay. For ubiquitination analysis, equal volumes of clarified lysates were divided, with one portion used directly for IB and the other subjected to IP, which was performed following the ChromoTek protocol with minor modifications. Briefly, antibody-conjugated agarose beads were equilibrated and incubated with lysates at 4°C overnight (16 h) with end-over-end rotation. For Myc-tagged proteins, anti-Myc agarose beads (30 μL per 500 μg protein; Cat. #yta, ChromoTek) were used; for mCherry-tagged proteins, anti-RFP agarose beads (50 μL per 500 μg protein; Cat. #rta, ChromoTek) were employed. After incubation, beads were washed with buffer containing 10 mM Tris/HCl (pH 7.5), 150 mM NaCl (Cat. #6001985, Fisher Scientific), 0.05% Nonidet P40 substitute (Cat. #11332473001, Sigma-Aldrich), and 0.5 mM EDTA (pH 7.2, 4°C; Cat. #E-5134, Sigma-Aldrich). Bound proteins were eluted in 2×SDS sample buffer (Cat. #C994D49, Thomas Scientific) by heating at 70°C for 10 min. Eluted samples were subjected to IB analyses to assess protein ubiquitination as previously described (63).

##### NT-CYP1A chimeras

Relative CYP1A templates were linearized with the corresponding primers (SI, **Table 1**). Then, homologous recombination was performed according to the NEB Gibson Assembly protocol (Manufacturer’s Instructions). Briefly, the fragment of 1A1 or 1A2 was mixed at a ratio of 3:1 with a total mass not exceeding 0.5 pmol in a 10 μL-volume. Subsequently, Gibson Assembly Master Mix (2X; 10 μL) was added, and the mixture was incubated in a thermocycler at 50°C for 30 min. The mixture (2 μL) was then transfected into competent *E. coli* cells.

##### NT-construct and tag replacement by site-directed mutagenesis

Site-directed mutagenesis was carried out using the Q5 Hot Start High-Fidelity 2× Master Mix (Cat. #E0554S, New England Biolabs) according to the manufacturer’s instructions. Briefly, PCR reactions (25 μL) contained 12.5 μL Master Mix, 0.5 μM forward and reverse primers, 1 ng DNA, and nuclease-free water, with cycling conditions of 98°C for 30 s, 25 cycles of 98°C for 10 s, 50–72°C for 10–30 s, and 72°C for 20–30 s/kb, followed by 72°C for 2 min. Amplified products (1 μL) were treated in a 10 μL Kinase, Ligase and DpnI enzymes mix (KLD) reaction mixture (New England Biolabs) for 5 min at room temperature and 5 μL of the mix was transformed into chemically competent cells (50 μL) for transformation and culture.

##### Plasmid construction for live imaging

To verify protein interactions in living cells, full-length (FL) or truncated rabbit CYP1A1, CYP1A2 or human ERLN1, ERLN2 and ODC1 coding sequences were tagged with cSURF and nSURF (Una-G)-tags respectively (by Gibson Assembly or Site-Directed Mutation) and cloned into pcDNA3.1 to create SURF-based protein-protein interaction (PPI) reporters (the SURF vector backbone was kindly provided by the Shu Lab, UCSF (56). All constructs were generated using standard enzyme digestion and ligation methods.

##### Preliminary soluble “S” vs pellet “P” fractionation

HepG2 cells were plated in a 15-cm^2^ plates, with 25 mL MEM, before plasmid transfection. Forty-eight h post-transfection, harvested cells were resuspended in a hypotonic buffer containing 10 mM 4-(2-Hydroxyethyl)-1-Piperazine ethanesulfonic Acid (HEPES, #BP310-1; Fisher Scientific), 10 mM KCl (#C752X06; Thomas Scientific), 1.5 mM MgCl_2_ (# AA12315A1; Fisher Scientific), 0.5 mM DTT (# C995K75; Thomas Scientific), and the protease inhibitor mixture, pH 7.4 and placed on ice for 30 min (29, 30, 70). The cells were lysed with 10 strokes in a glass homogenizer and then sonicated by the Omni Sonic Ruptor 250 sonicator at an amplitude level of 20 (50W) for 3 cycles, each consisting of five pulses with a minimum interval of one min between cycles. Unbroken cells and nuclei were removed by centrifugation at 600 g for 5 min. The supernatant was diluted to 0.5 mL with a membrane solubilization buffer containing 50 mM HEPES, pH 7.25, 150 mM NaCl, 5 mM EDTA, the protease inhibitor mixture, and Triton X-100 (# 97063-996; VWR) to a final concentration of 0.1 % v/v Triton X-100, and incubated on ice for 30 min. Samples were centrifuged using a TLA 120.1 rotor at 100,000 g for 1 h in a Beckmann CTX ultracentrifuge at 4°C. Supernatants (0.5 mL) were saved, and the pellets were resuspended in 0.5 mL of TISO buffer containing 8 M urea (Cat. #BP169-212, Fisher Scientific), 2 M thiourea (Cat. #62-56-6, Sigma-Aldrich), 4% CHAPS (Cat. #C802K33, Thomas Scientific), 20 mM Tris/HCl (Cat. #104028-344, VWR), and 30 mM dithiothreitol (DTT; Cat. #C995K73, Thomas Scientific) along with the protein inhibitor mixture. Protein distribution in the pellet (“P”) and supernatant (“S”) was then analyzed by IB analyses.

##### Isolation of mitochondria-associated endoplasmic reticulum (ER) membranes (MAMs) and detergent-resistant membranes (DRMs)

HepG2 cells were plated in a 15-cm^2^ plate, with 25 mL MEM, before plasmid transfection. Forty-eight h post-transfection, cells were harvested and resuspended in PBS (4.5 mL). Cells were aliquoted for MAM isolation (2 mL), DRM isolation (2 mL), and whole cell lysates (0.5 mL). For the MAM isolation, cells were resuspended in ice-cold MAM resuspension buffer, containing 225 mM mannitol (#17311-1KG-F; Sigma-Aldrich), 75 mM sucrose (#C993F64; Thomas Scientific), 0.1 mM EGTA (#C994Y58; Thomas Scientific), and 30 mM Tris-HCl, pH 7.4. The resuspended cells were then homogenized gently and slowly to break >80% cells. The homogenized extracts were then subjected to centrifugation at 600 g for 5 min at 4°C. The centrifugation was repeated and the supernatant collected only when no pellet traces were visible. The collected supernatants were transferred into 1.5 mL centrifuge tubes and centrifuged at 7,000 g with Eppendorf centrifuge 5415C for 10 min at 4°C to separate the crude mitochondrial pellet from the supernatant containing the cytosol and ER. The centrifugation was repeated and the supernatant collected only when no pellet traces remained to avoid contamination with MAMs and mitochondria. The collected supernatant was further ultracentrifuged at 100,000×g in a SW41Ti rotor for 1 h in an Optima XE-100 ultracentrifuge to isolate the ER (pellet) and cytoplasm (supernatant). The crude mitochondrial pellet was washed by IB cells-2 buffer [mannitol (225 mM), sucrose (75 mM), Tris/HCl (30 mM)] twice, then gently resuspended in 2 mL ice-cold mitochondrial resuspension buffer (MRB) containing mannitol (250 mM), HEPES (5 mM), and EGTA (0.5 mM), pH 7.4]. This suspension was then added on top of a Percoll medium, which included 8 mL of mannitol (250 mM), HEPES (5 mM, pH 7.4), and EGTA (0.5 mM) and 30% (v/v) Percoll (P4937-25ML), in a Seton open-top polyclear tube (#PN7030; Seton Scientific). Then, the MRB was carefully layered on top of the mitochondrial suspension to fill the centrifuge tube. The tube was then subjected to ultracentrifugation (in an Optima XE-100 ultracentrifuge) at 95,000 g in a SW41Ti Rotor for 60 min at 4 °C to separate the crude MAMs (top fraction) and pure mitochondria (bottom fraction). To obtain pure MAMs, the crude MAMs were diluted with MRB and centrifuged at 6300 g for 10 min at 4°C. The supernatant was then further ultracentrifuged at 100,000 g in a SW41Ti rotor for 1 h, and the resulting pellet was collected as the “purified” MAMs.

For the DRM isolation, cells were resuspended in a lysis buffer (50 mM HEPES, 150 mM NaCl, 5 mM EDTA, and the protease inhibitor mixture, pH 7.4) with 1% Triton X-100 to a final concentration of 0.1% Triton X-100 (v/v). The cells were lysed with 50 strokes of a homogenizer and following sonication with the Omni Sonic Ruptor 250 sonicator at an amplitude level of 20 (50 W) for 3 cycles in ice, each consisting of five pulses with a minimum interval of one min between cycles. Unbroken cells and nuclei were removed (pellet fraction) by centrifugation at 1000 g for 5 min at 4 °C. DRMs were isolated as described previously (71). In brief, sucrose solutions were prepared in a buffer of HEPES (50 mM), NaCl (150 mM), and the protease inhibitor mixture, pH 7.4. Solubilized samples were then combined with an equal volume of 80% sucrose solutions and placed at the bottom of a centrifuge tube. A discontinuous gradient was laid on top consisting of 6 ml of a 38% sucrose solution and 3 ml of a 5% sucrose solution. Samples were centrifuged for 19 h at 210,000 g at 4°C in a SW41Ti rotor in a Beckmann ultracentrifuge. Ten 1 mL-fractions were removed from the top of the gradient, and the pellet was homogenized in TISO buffer (1 mL). The protein distribution in the gradient was analyzed by IB analyses. DRMs float at the 5/38% interface (fractions 1–6) of the gradient.

To “wash” and only pool the main membrane fractions, DRM fractions were ultra-centrifuged at 105,000 g for 1 h at 4°C in a SW41Ti rotor in a Beckmann ultracentrifuge. The pellet was then homogenized with 1.15% ice-cold KCl. This wash step was repeated twice, and the final ultracentrifuged pellet was resuspended in TISO buffer with a protease inhibitor mixture and employed as the “DRM-fraction”. Sucrose-gradient fractions 7-10 were considered as “non-DRMs”, and similarly diluted, similarly “washed” with 1.15% KCl and resedimented by ultracentrifugation at 105,000 g for 1 h at 4°C in a SW41Ti-rotor in a Beckmann ultracentrifuge.

##### Densitometirc Scanning Analyses

The densitometric scanning analyses were performed by ImageJ software. The density of each band in the same gel was analyzed and the densitometric target protein/internal reference protein ratios determined.

##### Erlin-1 knockdown and rescue

Predesigned siRNA duplexes specific to human erlin mRNAs were procured from Qiagen. For the knockdown of erlin-1, two erlin-1-specific siRNAs were utilized: Hs_SPFH1_2, (cat. #SI00731402; Qiagen), and Hs_SPFH1_3, (cat. #SI00731409; Qiagen), designated as siRNA 1 and siRNA 2 (siRNA1/2) respectively. The "AllStars negative control siRNA" (Cat. #SI03650318; Qiagen) served as the negative (scrambled siRNA) control. HepG2 cells were transfected with siRNA using the HiPerFect Transfection Reagent (Cat. #301705; Qiagen). The cells were transfected in 6-well plates with a volume of 2 mL/well. Cells were harvested 72 h post-transfection, after preliminary evaluation of maximal knockdown (KD). Cells were homogenized in RIPA buffer with the protease-inhibitor mixture and sonicated using a microsonic cell disruptor at an amplitude of 20 (50W) for three cycles, each cycle comprising of five pulses with a minimum one-min interval between cycles. Unbroken cells and nuclei were removed (pellet fraction) by centrifugation at 600 g for 5 min at 4°C. To assess the cellular aggregation following erlin-1 KD, the homogenates were centrifuged at 14,000 x g for 10 min at 4°C in a Eppendorf Centrifuge 5425, and the supernatant was used as the soluble fraction. The pellet obtained from this centrifugation was solubilized with the same volume of TISO buffer containing the protease inhibitor mixture and employed as the “pellet” fraction.

To rescue erlin-1-KD, we first constructed two plasmids (**SI**, **Table 1**): i) erlin-1 (DNA mutants), which contained several nucleotide mutations in its DNA sequence, while retaining the corresponding native amino acid sequence, and ii) erlin-1 N(1–30)-mCherry, in which the NT 1-30 residue-subdomain of erlin-1 is fused to mCherry. Subsequently, 12 h after addition of siRNAs 1/2, the medium was replaced, and plasmids containing siRNAs 1/2 and either erlin-1 (DNA mutants) or erlin-1 N(1–30)-mCherry were added. After 24 h, the medium was replaced with fresh MEM containing siRNA 1/2, and the cells were further cultured for another 24 h. At this stage (72 h total), the cells were in a state of being rescued after erlin-1-KD and were ready for use in further assays.

##### Immunoblotting (IB) analyses

Protein concentrations were determined by the BCA assay and equal amounts of proteins (5 μg) were separated on SDS-PAGE gels (4-15%: cat. #3450029; Bio-Rad Laboratories, and 7.5%: cat. #5671025; Bio-Rad Laboratories).

Proteins were transferred onto 0.2 μm nitrocellulose membranes (cat. #162011; Bio-Rad Laboratories) for IB analyses at 4°C for 1 h. Commercial primary antibodies, including mCherry (cat. #26765-1-AP; Proteintech), ATG5 polyclonal antibody (ATG5, cat. #10181-2-AP; Proteintech), Mitofusin-2 (D1E9) rabbit mAb (MFN2, cat. #11925S, Cell Signaling Technology), VDAC1/Porin Antibody (B-6) (VDAC1, cat. #sc-390996; Santa Cruz Biotechnology), erlin-1 (cat. #17311-1-AP; Proteintech Group), P450 Reductase (CPR, cat. #29814-1-AP; Proteintech Group), Myc-Tag (cat. #9402S; Cell Signaling Technology), ubiquitin (cat. #10201-2-AP; Proteintech Group), LC3A/B (cat. #12741T; Cell Signaling Technology), were used for detecting the primary protein. GAPDH (cat. #10494-1-AP; Proteintech) or Histone-H3 (cat. #ab1791-100UG; Abcam) were used as the internal reference proteins (loading controls). The secondary antibodies were horse-radish peroxidase (HRP)-linked anti-rabbit antibody (cat. #7074S; Cell Signaling Technology) or HRP-linked anti-mouse antibody (cat. #7076S; Cell Signaling Technology), respectively.

##### U2OS transfections for imaging

Before cell culture, glass coverslips (Cat. #12-541B, Thermo-Fisher Scientific) were placed in 35-mm culture dishes (Cat. #08-774-258, Thermo-Fisher Scientific), rinsed twice with 2 mL phosphate-buffered saline (PBS, pH 7.2; Cat. #10010002, Gibco), air-dried, and sterilized under ultraviolet light for 1 h. Each coverslip was then covered with 2 mL McCoy’s 5A modified medium (Cat. #16600108, Gibco) supplemented with 10% FBS and antibiotics. U2OS cells were seeded at a density of 1 × 10⁵ cells/mL (1 mL per dish) and cultured under standard conditions (37°C, 5% CO_2_/95% O_2_) for 24 h. Transfection was carried out using the Lipofectamine 3000 transfection reagent (Cat. #L3000015, Thermo-Fisher Scientific) with 1 μg plasmid DNA per dish, following the manufacturer’s protocol. After 12 h, the medium was replaced with fresh complete medium, and cells were further cultured for at least 24 h before fixation with 4% formaldehyde (Cat. #28908, Thermo Scientific).

##### Intracellular colocalization of CYP1A2-mCherry and erlin-1 by confocal imaging

U2OS cells cultured as described earlier were transfected with CYP1A2-mCherry (1 μg/well) in 6-well plates. After 48 h-transfection, cells were washed with cold PBS once, and then fixed with 4% formaldehyde for 20 min at room temperature followed by methanol at −20°C for 1 min. Cells were then rinsed with PBS 3 times for 5 min each and treated with 0.4% Triton X-100 in PBS for 10 min for cell permeabilization. The cells were rinsed twice in PBS for 5 min each to wash off the Triton X-100. Cells were blocked for 1 h with 10% normal goat serum in PBS/0.1% Tween-20 at room temperature, and then stained with erlin-1 polyclonal antibody (Cat #17311-1-AP, Proteintech) at 4°C overnight. Nuclei were visualized with blue fluorescent protein-Histone 2B (BFP-H2B, cat #55243, Addgene). Cells were then washed in PBS/0.1% Tween thrice and then stained with goat anti-rabbit IgG Alexa Fluor 488 (Invitrogen) for 1 h at room temperature. Cells were further washed thrice in PBS/0.1% Tween and then mounted using ProLong Diamond Anti-fade Mountant with DAPI nuclear stain (Molecular Probes, Grand Island, NY). Images were taken with a Nikon Yokogawa CSU-22 Spinning Disc Confocal Microscope using a Plan Apo VC 60 3/1.4 oil lens. All panels were subjected to identical exposure times.

##### The fluorogenic split SURF-bimolecular functional complementation assay

U2OS cells cultured as described above were transfected with the Lipofectamine 3000 transfection reagent and a total of 1 μg plasmid mix consisting of 400 ng cSURF fused protein, 400 ng nSURF fused protein, 200 ng eBFP2-H2B. HEK293T cells (ATCC, No. 3216) were cultured in DMEM (Dulbecco’s Modified Eagle Medium Gibco, cat. #11965-092) supplemented with 10% fetal bovine serum (FBS; Gibco, #16000044) and 1% penicillin Streptomycin (Gibco, cat. #15140-122) and seeded in an 8-chambered coverglass plate (Cellvis, cat. #C8-1.5P). Cells were incubated at 37°C in a humidified atmosphere under 5%CO_2_/95%O_2_. After reaching 40-50% confluency, plasmids were transfected into cells using calcium chloride. Briefly, a total of 250 ng plasmid mix consisting of 100 ng cSURF fused protein, 100 ng nSURF fused protein, and 50 ng eBFP2-H2B were mixed with 10 μL 1 x Hank’s Balanced Salts buffer (HBS) (HyClone, cat. #SH30588.02), 0.6 L of 2 M Calcium Chloride (cat. #C25100, RPI) added, mixed gently and incubated for 15-20 min at room temperature. The DNA/HBS complexes were then added into each cell-containing well, mixed gently by rocking the plate back and forth. The U2OS and HEK293T cells were incubated at 37°C in a humidified atmosphere of 5%CO_2_/95%O_2_ for 18-24 h. 10 M Chromophore Bilirubin (cat. #2011, Calbiochem) was then added and the cells further incubated for 15-20 min before live-cell imaging analyses. All live-cell imaging analyses were conducted using a Nikon Eclipse Ti inverted microscope, featuring a Yokogawa CSU-W1 confocal scanner unit (Andor), an ORCA-Flash 4.0 digital CMOS camera (Hamamatsu), an ASI MS-2000 XYZ automated stage, and a Nikon Plan Apo 60x oil objective lens (N-PLANAPO-60X). The setup also included a CO_2_/temperature control unit. Laser inputs were provided by an Integrated Laser Engine (Spectral Applied Research) with coherent laser lines at 488 nm for SURF imaging and 405 nm for BFP imaging. Images were captured with exposure times of 2 sec and acquisition was managed using NIS-Elements Microscope Imaging Software (Nikon). Image processing was performed with ImageJ software.

##### Statistical analyses

All data are presented as mean ± standard deviation (SD). Statistical significance was determined using an unpaired two-tailed Student’s t-test for comparisons between two groups, or one-way ANOVA for comparisons among more than two groups. A value of p < 0.05 was considered statistically significant (*) over that not statistically significant (ns). All statistical analyses were performed using GraphPad Prism 6.07.

## Acknowledgments

We sincerely thank Drs. Bo Huang and Klaus Yserentant (Dept. of Pharmaceutical Chemistry, UCSF) for their valuable input, helpful suggestions and some preliminary findings. We also thank the members of the UCSF Nikkon Center, for their valuable input and Dr. Xing Liang (Stanford University) for her valuable assistance with the U2OS cell confocal imaging and quantification.

