## Supporting Table & Figures for "Hepatic cytochrome P450 endoplasmic reticulum-associated degradation (ERAD): Topological determinants and cellular partnerships that dictate the preferential P450 proteolytic sorting into macroautophagy rather than UPS"

### X. Hong et al.

### SUPPORTING INFORMATION

### (Table 1 and Figs S1-S7)

**SI-Table 1**

| Construct | Template | Primer |
|---|---|---|
| CYP1A1-mCherry | pmCherry-N1 | 5'-CATGGTGAGCAAGGGCGAGGAG-3' |
| | | 5'-CATGGTGGCGTAGCGGATC-3' |
| | CYP1A1-EGFP | 5'-GATCGGCTAGCGGCACCACCATGCTTTTCCAAATCTCCATGTCGGC-3' |
| | | 5'-CTCCTCGGCCTTGCTCACCATG-3' |
| CYP1A2-mCherry | pmCherry-N1 | 5'-CATGGTGAGCAAGGGCGAGGAG-3' |
| | | 5'-CATGGTGGCGTAGCGGATC-3' |
| | CYP1A2-EGFP | 5'-GATCGGCTAGCGGCACCACCATGGCATTGTCCCAGCTCTGTTCCC-3' |
| | | 5'-CTCCTCGGCCTTGCTCACCATG-3' |
| CYP2B1-mCherry | pmCherry-N1 | 5'-CATGGTGAGCAAGGGCGAGGAG-3' |
| | | 5'-CATGGTGGCGTAGCGGATC-3' |
| | CYP2B1-Myc-His | 5'-GATCGGCTAGCGGCACCACCATGGCATTGTCCCAGCTCTGTTCCC-3' |
| | | 5'-CTCCTCGGCCTTGCTCACCATG-3' |
| CYP2B6-mCherry | pmCherry-N1 | 5'-CATGGTGAGCAAGGGCGAGGAG-3' |
| | | 5'-CATGGTGGCGTAGCGGATC-3' |
| | CYP2B6-Myc-His | 5'-GATCGGCTAGCGGCACCACCATGGCATTGTCCCAGCTCTGTTCCC-3' |
| | | 5'-CTCCTCGGCCTTGCTCACCATG-3' |
| 1A1(1–109)1A2-mCherry | CYP1A1-mCherry | 5'-AGATCCGCTAGCGCCACCATGCTTTTCCCAATCTCCATG-3' |
| | | 5'-AGGGTGGAGGTGTAGAGGTCaggccggcccttgaaatc-3' |
| | CYP1A2-mCherry | 5'-GGTGGCGCTAGCGGATCT-3' |
| | | 5'-GACCTCTACACCTCCACCCTC-3' |
| 1A2(1–107)1A1-mCherry | CYP1A1-mCherry | 5'-GGTGGCGCTAGCGGATCT-3' |
| | | 5'-GACCTCTACACCTTCACCCTC-3' |
| | CYP1A2-mCherry | 5'-GGTGAAGGTGTAGAGGTCGGGCCGGCCCTTGAAATC-3' |
| | | 5'-GAACCGTCAGATCCGCTAGCGCCACC-3' |
| 1A2(1-205)1A1-mCherry | CYP1A1-mCherry | 5'-GGTGGCGCTAGCGGATCT-3' |
| | | 5'-CGGCGCTATGACCACAAC-3' |
| | CYP1A2-mCherry | 5'-GTTGTGGTCATAGCGCCGtccgaagcacatggcacc-3' |
| | | 5'-AGATCCGCTAGCGCCACC-3' |
| 1A1-mCherry-Myc-His | CYP1A1-mCherry | 5'-aggatctgaatagcgccgtcgaccatcatcatcatcatcatTAGCGGCCGCGACTC-3' |
| | | 5'-cttctgagatgagtttttgttcgggcccaagcttggtaccCTTGTACAGCTCGTCCATGC-3' |

| | | |
|---|---|---|
| 1A2-mCherry-Myc-His | CYP1A2-mCherry | 5'-aggatctgaatagcgccgtcgaccatcatcatcatcatcatTAGCGGCCGCGACTC-3' |
| | | 5'-cttctgagatgagtttttgttcgggcccaagcttggtaccCTTGTACAGCTCGTCCATGC-3' |
| 1A1(1-109)1A2-mCherry-Myc-His | 1A1(1-109)1A2-mCherry | 5'-aggatctgaatagcgccgtcgaccatcatcatcatcatcatTAGCGGCCGCGACTC-3' |
| | | 5'-cttctgagatgagtttttgttcgggcccaagcttggtaccCTTGTACAGCTCGTCCATGC-3' |
| 1A2(1-107)1A1-mCherry-Myc-His | 1A2(1-107)1A1-mCherry | 5'-aggatctgaatagcgccgtcgaccatcatcatcatcatcatTAGCGGCCGCGACTC-3' |
| | | 5'-cttctgagatgagtttttgttcgggcccaagcttggtaccCTTGTACAGCTCGTCCATGC-3' |
| 1A2(1-205)1A1-mCherry -Myc-His | 1A2(1-205)1A1-mCherry | 5'-aggatctgaatagcgccgtcgaccatcatcatcatcatcatTAGCGGCCGCGACTC-3' |
| | | 5'-cttctgagatgagtttttgttcgggcccaagcttggtaccCTTGTACAGCTCGTCCATGC-3' |
| 2B1-Myc | 2B1-Myc-His | 5'-CAGATCCTCTTCTGAGATG-3' |
| | | 5'-tgatgaTGAGTTTAAACCCGCTG-3' |
| 2B6-Myc | 2B6-Myc-His | 5'-tgatgaTGAGTTTAAACCCGCTG-3' |
| | | 5'-CAGATCCTCTTCTGAGATG-3' |
| 2B1(N:1-30)-mCherry-Myc-His | 2B1-Myc-His | 5'-ATCCTCCTCGCCCTTGCTCACGGGCCCAAGCTTGGTACCgaagttgccacgggactttgg-3' |
| | | 5'-GAACAAAAACTCATCTCAGAAGAGGAT-3' |
| | CYP1A2-mCherry | 5'-GTGAGCAAGGGCGAGGAGGAT-3' |
| | | 5'-gctattcagatcctcttctgagatgagtttttgttc-3' |
| 2B6(N:1-30)-mCherry-Myc-His | 2B6-Myc-His | 5'-ATCCTCCTCGCCCTTGCTCACGGGCCCAAGCTTGGTACCgaggcggtcatgggtg-3' |
| | | 5'-GAACAAAAACTCATCTCAGAAGAGGAT-3' |
| | CYP1A2-mCherry | 5'-GTGAGCAAGGGCGAGGAGGAT-3' |
| | | 5'-gctattcagatcctcttctgagatgagtttttgttc-3' |
| 1A1(N:1–33)-mCherry | CYP1A1-mCherry | 5'-GATCCACCGGTCGCCACC-3' |
| | | 5'-TGAGGCCCTGATTACCCAG-3' |
| 1A2(N:1–33)-mCherry | CYP1A2-mCherry | 5'-GGATCCCCACCGGTCGCC-3' |
| | | 5'-CAAACCCTTGAGCACCCAGAATACC-3' |
| 1A2-nSURF | CYP1A2-mCherry | 5'-tccgttgatggagaagcgcag-3' |
| | | 5'-CGGCCGCGACTCTAGATCATA-3' |
| | SURF Plasmid | 5'-ctgcgcttctccatcaacggaaaaaaagcgccagtgctccagaaattcgtagga-3' |
| | | 5'-TATGATCTAGAGTCGCGGCCGgcgCTAgcgacgcccgtcagaaggaaactc-3' |
| 1A2-cSURF | CYP1A2- | 5'-tccgttgatggagaagcgcag-3' |

| | | |
|---|---|---|
| | mCherry | 5'-CGGCCGCGACTCTAGATCATA-3' |
| | SURF Plasmid | 5'-ctgcgcttctccatcaacggaaaaaaagcgGGTGTTAAAAGCGTCGTTAACCTG-3' |
| | | 5'-TATGATCTAGAGTCGCGGCCGgcgCTAgcgctccgtcgccctgcgata-3' |
| 1A1-nSURF | CYP1A1-mCherry | 5’- AGAGCGCAGCTGCATTTGGAA-3’ |
| | | 5'-TAGCGGCCGCGACTCTAGATC-3' |
| | SURF Plasmid | 5’-GATCTAGAGTCGCGGCCGCTAacgcccgtcagaaggaaactc-3’ |
| | | 5'-TTCCAAATGCAGCTGCGCTCTccagtgctccagaaattcgtagga-3' |
| 1A1-cSURF | CYP1A1-mCherry | 5’- AGAGCGCAGCTGCATTTGGAA-3’ |
| | | 5'-TAGCGGCCGCGACTCTAGATC-3' |
| | SURF Plasmid | 5’-GATCTAGAGTCGCGGCCGCTAggtgttaaaagcgtcgttaacctg-3’ |
| | | 5'-TTCCAAATGCAGCTGCGCTCTctccgtcgccctgcgata-3' |
| 1A1(N:1-33)-nSURF | CYP1A1-mCherry | 5'-gagtttccttctgacgggcgtTAGCGGCCGCGACTCTAG-3' |
| | | 5'-agttcctacgaatttctggagcacaaagggTGAGGCCCTGATTACCCA-3' |
| | SURF Plasmid | 5'-gtgctccagaaattcgtaggaact-3' |
| | | 5'-acgcccgtcagaaggaaactc-3' |
| 1A1(N:1-33)-cSURF | CYP1A1-mCherry | 5'-tatcgcagggcgacggagTAGCGGCCGCGACTCTAG-3' |
| | | 5'-CAGGTTAACGACGCTTTTAACACCaaagggTGAGGCCCTGATTACCCA-3' |
| | SURF Plasmid | 5'-GGTGTTAAAAGCGTCGTTAACCTG-3' |
| | | 5'-ctccgtcgccctgcgata-3' |
| 1A2(N:1-33)-nSURF | CYP1A2-mCherry | 5'-gagtttccttctgacgggcgtTAGCGGCCGCGACTCTAG-3' |
| | | 5'-agttcctacgaatttctggagcacaaagggcaaacccttgagcacccagaa-3' |
| | SURF Plasmid | 5'-gtgctccagaaattcgtaggaact-3' |
| | | 5'-acgcccgtcagaaggaaactc-3' |
| 1A2(N:1-33)-cSURF | 1A2-cSURF | 5'-GGAAAAAAAGCGGGTG-3' |
| | | 5'-CAAACCCTTGAGCAC-3' |
| Erlin-1-mCherry | CYP1A2-mCherry | 5'-CATGGTGAGCAAGGGCGAGGAG-3' |
| | | 5'-CATGGTGGCGTAGCGGATC-3' |
| | Erlin-1-FLAG | 5'-CCGCTAGCGCCACCATGAATATGACTCAAGCCCGG-3' |
| | | 5'-GACCGGTGGGGATCCTGTGCTCTCTTTGTTTTGGA |

| | | |
|---|---|---|
| | | T-3' |
| Erlin-1(DNA Mutation)-mCherry | Erlin-1-mCherry | 5'-cctcgaactcaaAAGTACCAGGCCATTG-3' |
| | | 5'-tactcgggagtCAACTTGTGCTTGTTTG-3' |
| Erlin-2-mCherry | Erlin-2-Myc-FLAG | 5'-GCGATCGCCATGGCTCAG-3' |
| | | 5'-TGTGGGTGCCTCGAGGGG-3' |
| | CYP1A2-mCherry | 5'-CCCCTCGAGGCACCCACAGGATCCCCACCGGTCG-3' |
| | | 5'-CTGAGCCATGGCGATCGCCATGGTGGCGCTAGCG-3' |
| Erlin-1-nSURF | Erlin-1-mCherry | 5'-tagcagacttcctctgccctcTGTGCTCTCTTTGTTTTGGAT-3' |
| | | 5'-ATGAATATGACTCAAGCCCGG-3' |
| | 1A2-nSURF | 5'-gagggcagaggaagtctgctaGGAAAAAAAGCGCCAGTG-3' |
| | | 5'-ccgggcttgagtcatattcatGGTGGCGCTAGCGG-3' |
| Erlin-1-cSURF | Erlin-1-mCherry | 5'-GACCGGTGGGGATCCTGTGCTCTCTTTGTTTTGGAT-3' |
| | | 5'-TAAACTCGAGTCTAGAGCGGC-3' |
| | SURF Plasmid | 5'-AGCACAGGATCCCCACCGGTCggtgttaaaagcgtcgttaacctg-3' |
| | | 5'-GCCGCTCTAGACTCGAGTTTActccgtcgccctgcgata-3' |
| Erlin-2-nSURF | Erlin-2-Myc-FLAG | 5'-gagtttccttctgacgggcgtGATGACGACGATAAGGTT-3' |
| | | 5'-agttcctacgaatttctggagcacATCCAGGATATCATTTGC-3' |
| | 1A2-nSURF | 5'-gtgctccagaaattcgtaggaact-3' |
| | | 5'-acgcccgtcagaaggaaactc-3' |
| Erlin-2-cSURF | Erlin-2-Myc-FLAG | 5'-GCGATCGCCATGGCTCAGTTG-3' |
| | | 5'-GATGAGTTTCTGCTCGAGCGG-3' |
| | 1A2-cSURF | 5'-CCGCTCGAGCAGAAACTCATCGCGGGTGTTAAAAGCGTCGTT-3' |
| | | 5'-CAACTGAGCCATGGCGATCGCGGTGGCGCTAGCGGATCT-3' |
| Erlin-2(T65I)-cSURF | Erlin-2-cSURF | 5'-TGTACAGACCatcCTCCAAACTG-3' |
| | | 5'-GACTTATAGGATGTGATGAAC-3' |
| Erlin-1(N:1-30)-nSURF | Erlin-1-nSURF | 5'-GAGGGCAGAGGAAGTC-3' |
| | | 5'-CTCCTCAATCTTGTGGATG-3' |
| Erlin-1(N:1-30)-cSURF | Erlin-1-cSURF | 5'-CTCCTCAATCTTGTGGATGGAGG-3' |
| | | 5'-GGATCCCCACCGGTCG-3' |
| Erlin-2(N:1-30)-nSURF | Erlin-2-nSURF | 5'-AAGGAGAACACGCGTAC-3' |
| | | 5'-ATGTCCCTCTTCTATCTTGTG-3' |
| Erlin-2(N:1-30)-cSURF | Erlin-2-cSURF | 5'-AAGGAGAACACGCGTAC-3' |
| | | 5'-ATGTCCCTCTTCTATCTTGTG-3' |
| ODC-cSURF | pET28a-His6- | 5'-atgggcagcagccatcatcat-3' |

| | | |
|---|---|---|
| | ODC-ctag | 5'-GACGCTTTTAACACCcgcttttttccagcctctggctcaccgct-3' |
| | 1A2-cSURF | 5'-ggaaaaaaagcgGGTGTTAAAAGCGTC-3' |
| | | 5'-atgatgatggctgctgcccatGGTGGCGCTAGCGG-3' |
| 1A2-iRFP | 1A2-mCherry | 5'-atcggcgtgatggaagagTAGCGGCCGCGACTC-3' |
| | | 5'-cgacggatccttcagccatTCCGTTGATGGAGAAGCGC-3' |
| | iRFP Plasmid | 5'-atggctgaaggatccgtcg-3' |
| | | 5'-ctcttccatcacgccgat-3' |
| Erlin-1-iRFP | Erlin-1-FLAG | 5'-TGTGCTCTCTTTGTTTTGGA-3' |
| | | 5'-TAAACTCGAGTCTAGAGCGG-3' |
| | iRFP Plasmid | 5'-TCCAAAACAAAGAGAGCACAatggctgaaggatccgtcg-3' |
| | | 5'-CCGCTCTAGACTCGAGTTTActcttccatcacgccgat-3' |
| Erlin-1(N:1-30)-iRFP | Erlin-1-iRFP | 5'-GGTGGGGGTGGAGGC-3' |
| | | 5'-CTCCTCAATCTTGTGGATGGAGG-3' |

**SUPPORTING INFORMATION: Figures SF1-SF7**

Fig. S1

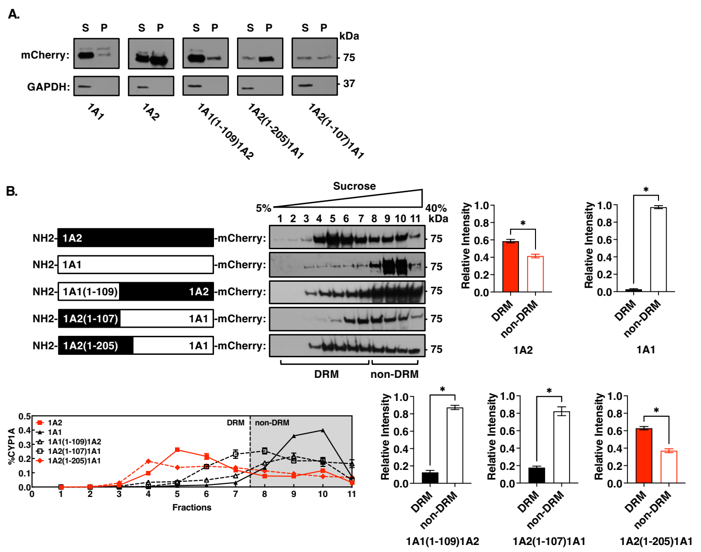
A.
S P
mCherry:
GAPDH:
kDa
75
37
1A1
1A2
1A1(1-109)1A2
1A2(1-205)1A1
1A2(1-107)1A1
B.
Sucrose
5%
40%
1 2 3 4 5 6 7 8 9 10 11
kDa
NH2- 1A2 -mCherry:
NH2- 1A1 -mCherry:
NH2- 1A1(1-109) 1A2 -mCherry:
NH2- 1A2(1-107) 1A1 -mCherry:
NH2- 1A2(1-205) 1A1 -mCherry:
DRM
non-DRM
Relative Intensity
%CYP1A
Fractions
1A2
1A1
1A1(1-109)1A2
1A2(1-107)1A1
1A2(1-205)1A1
1A2(1-205)1A1

Fig. S2

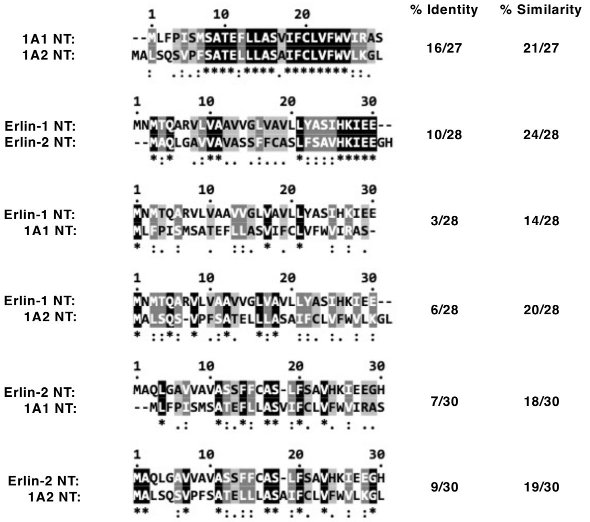
% Identity
% Similarity
1A1 NT: --MLFPISMSATEFLLASVIFCLVFWVIRAS
1A2 NT: MALSQSVPFSATELLLASAIFCLVFWVLKGL
16/27
21/27
Erlin-1 NT: MNMTQARVLVAAVVGLVAVLLYASIHKIEE--
Erlin-2 NT: --MAQLGAVVAVASSFFCASLFSAVHKIEEGH
10/28
24/28
Erlin-1 NT: MNMTQARVLVAAVVGLVAVLLYASIHKIEE
1A1 NT: MLFPISMSATEFLLASVIFCLVFWVIRAS-
3/28
14/28
Erlin-1 NT: MNMTQARVLVAAVVGLVAVLLYASIHKIEE--
1A2 NT: MALSQS-VPFSATELLLASAIFCLVFWVLKGL
6/28
20/28
Erlin-2 NT: MAQLGAVVAVASSFFCAS-LFSAVHKIEEGH
1A1 NT: --MLFPISMSATEFLLASVIFCLVFWVIRAS
7/30
18/30
Erlin-2 NT: MAQLGAVVAVASSFFCAS-LFSAVHKIEEGH
1A2 NT: MALSQSVPFSATELLLASAIFCLVFWVLKGL
9/30
19/30

**Fig.S3**

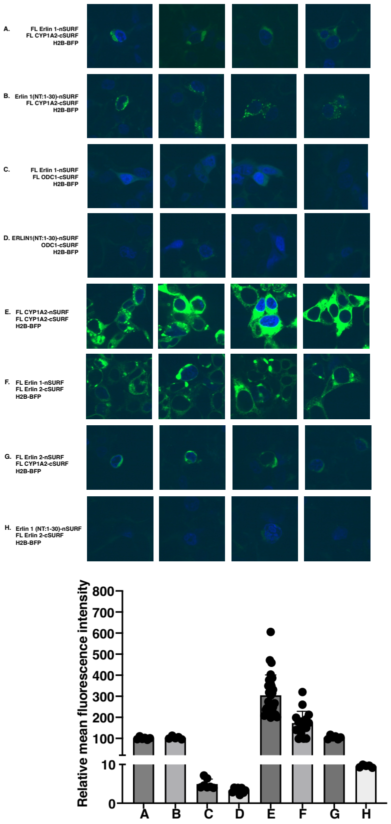
A.
FL Erlin 1-nSURF
FL CYP1A2-cSURF
H2B-BFP
B.
Erlin 1(NT:1-30)-nSURF
FL CYP1A2-cSURF
H2B-BFP
C.
FL Erlin 1-nSURF
FL ODC1-cSURF
H2B-BFP
D.
ERLIN1(NT:1-30)-nSURF
ODC1-cSURF
H2B-BFP
E.
FL CYP1A2-nSURF
FL CYP1A2-cSURF
H2B-BFP
F.
FL Erlin 1-nSURF
FL Erlin 2-cSURF
H2B-BFP
G.
FL Erlin 2-nSURF
FL CYP1A2-cSURF
H2B-BFP
H.
Erlin 1 (NT:1-30)-nSURF
FL Erlin 2-cSURF
H2B-BFP
Relative mean fluorescence intensity
800
700
600
500
400
300
200
100
10
0
A
B
C
D
E
F
G
H

#### Fig. S4

A.

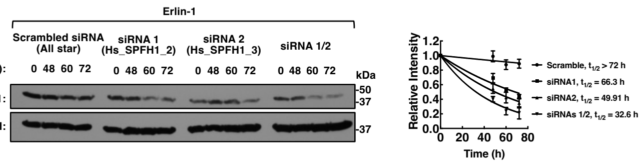
Erlin-1
Scrambled siRNA (All star)
siRNA 1 (Hs_SPFH1_2)
siRNA 2 (Hs_SPFH1_3)
siRNA 1/2
Time (h):
0 48 60 72
0 48 60 72
0 48 60 72
0 48 60 72
kDa
Erlin-1:
-50
-37
GAPDH:
-37
Relative Intensity
1.2
1.0
0.8
0.6
0.4
0.2
0.0
0 20 40 60 80
Time (h)
Scramble, $t_{1/2}$ > 72 h
siRNA1, $t_{1/2}$ = 66.3 h
siRNA2, $t_{1/2}$ = 49.91 h
siRNAs 1/2, $t_{1/2}$ = 32.6 h

**Fig. S5**

**A.**

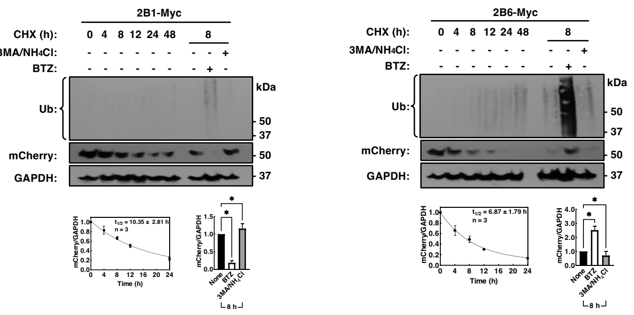
2B1-Myc
CHX (h): 0 4 8 12 24 48 8
3MA/NH4Cl: - - - - - - - - +
BTZ: - - - - - - - + -
Ub:
kDa
50
37
mCherry:
50
GAPDH:
37
t1/2 = 10.35 ± 2.81 h
n = 3
mCherry/GAPDH
Time (h)
None
BTZ
3MA/NH4Cl
8 h
2B6-Myc
CHX (h): 0 4 8 12 24 48 8
3MA/NH4Cl: - - - - - - - - +
BTZ: - - - - - - - + -
Ub:
kDa
50
37
mCherry:
50
GAPDH:
37
t1/2 = 6.87 ± 1.79 h
n = 3
mCherry/GAPDH
Time (h)
None
BTZ
3MA/NH4Cl
8 h

**B.**

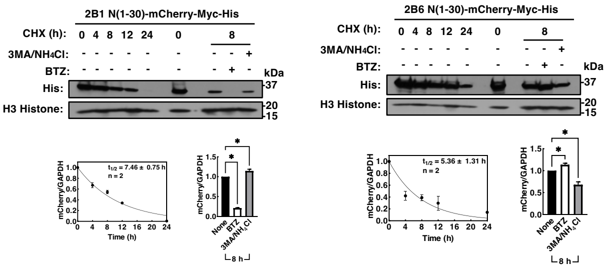
2B1 N(1-30)-mCherry-Myc-His
CHX (h): 0 4 8 12 24 0 8
3MA/NH4Cl: - - - - - - - - +
BTZ: - - - - - - - + -
kDa
His:
37
H3 Histone:
20
15
t1/2 = 7.46 ± 0.75 h
n = 2
mCherry/GAPDH
Time (h)
None
BTZ
3MA/NH4Cl
8 h
2B6 N(1-30)-mCherry-Myc-His
CHX (h): 0 4 8 12 24 0 8
3MA/NH4Cl: - - - - - - - - +
BTZ: - - - - - - - + -
kDa
His:
37
H3 Histone:
20
15
t1/2 = 5.36 ± 1.31 h
n = 2
mCherry/GAPDH
Time (h)
None
BTZ
3MA/NH4Cl
8 h

#### Fig. S6

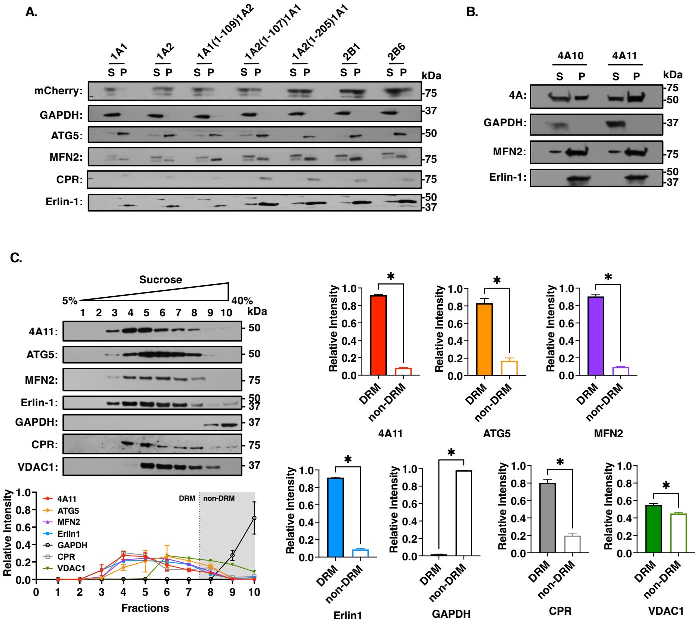

A.
1A1
1A2
1A1(1-109)1A2
1A2(1-107)1A1
1A2(1-205)1A1
2B1
2B6
S P
kDa
mCherry:
GAPDH:
ATG5:
MFN2:
CPR:
Erlin-1:
75
37
50
75
75
50
37
B.
4A10
4A11
4A:
GAPDH:
MFN2:
Erlin-1:
C.
Sucrose
5%
40%
1 2 3 4 5 6 7 8 9 10
4A11:
ATG5:
MFN2:
Erlin-1:
GAPDH:
CPR:
VDAC1:
Relative Intensity
Fractions
DRM
non-DRM
4A11
ATG5
MFN2
Erlin1
GAPDH
CPR
VDAC1
*

Fig.S7

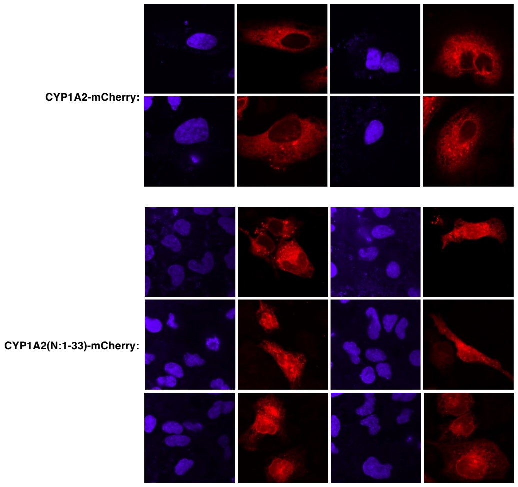
